# Language Models Learn Sentiment and Substance from 11,000 Psychoactive Experiences

**DOI:** 10.1101/2022.06.02.494544

**Authors:** Sam Freesun Friedman, Galen Ballentine

## Abstract

With novel hallucinogens poised to enter psychiatry, we lack a unified framework for quantifying which changes in consciousness are optimal for treatment. Using transformers (i.e. BERT) and 11,816 publicly-available drug testimonials, we first predicted 28-dimensions of sentiment across each narrative, validated with psychiatrist annotations. Secondly, BERT was trained to predict biochemical and demographic information from testimonials. Thirdly, canonical correlation analysis (CCA) linked 52 drugs’ receptor affinities with testimonial word usage, revealing 11 latent receptor-experience factors, mapped to a 3D cortical atlas. Together, these 3 machine learning methods elucidate a neurobiologically-informed, temporally-sensitive portrait of drug-induced subjective experiences. Different models’ results converged, revealing a pervasive distinction between lucid and mundane phenomena. MDMA was linked to “Love”, DMT and 5-MeO-DMT to “Mystical Experiences”, and other tryptamines to “Surprise”, “Curiosity” and “Realization”. Applying these models to real-time biofeedback, practitioners could harness them to guide the course of therapeutic sessions.

## Introduction

Quantifying the diversity, subtlety and trajectory of human feeling is difficult. However, reliably measuring these rich spectra of emotion is critical for the burgeoning field of psychedelic-assisted therapy, where the subjective qualities of drug-induced “trips” have repeatedly been shown to correlate with clinical efficacy [1–3]. For example, acute experiences involving overwhelming feelings of love, ego-dissolution, and mysticism correlate positively with long-term decreases in symptoms of anxiety and depression [4]. Relatedly, conventional psychiatric medications like antidepressants and antipsychotics target distinct but overlapping subjective experiences like anhedonia, paranoia, anxiety, and hallucinations. As novel molecules are introduced into standard psychiatric practice it is essential to establish a unifying paradigm–a rosetta stone–which translates how drugs modulate both neurochemistry and subjective experience [5]. Machine learning can assist in this translation by revealing latent structures that link phenomenal and biochemical levels of drug experience across the wide range of psychiatric, recreational, and psychedelic drug classes [6].

While emotions are fundamental to human affairs, scientists still lack a consistent neurobiological basis to describe and quantify them [7]. While gross aspects like arousal and hedonic tone are conserved across our species, the diverse subjective manifestations of psychiatric illnesses and the characteristic intensification of emotion and altered self-perception during acute psychedelic [8] experiences require more nuanced, multi-dimensional classifications [9]. Recent advances in Natural Language Processing (NLP) have operationalized this ambiguity by providing taxonomies of emotion derived from manually-annotated datasets of text [10–12]. Simultaneously, transformer-based models have made progress on many language tasks, and are increasingly able to learn generally applicable representations [13–15]. Here, we use models pre-trained on sentiments found in movie reviews and Reddit posts to estimate and disentangle emotional affect described in testimonials from both acute and chronic drug experiences [10].

Feelings unfold over time. This temporal trajectory of sentiment is of great significance; it is literally the substrate in which significance is felt. Moreover, the form of a sentiment’s trajectory shapes the memory that is ultimately consolidated. For example, the “peak-end rule” states that the intensities of the peak and the end of an experience determine its imprint in memory, while the notion of “primacy” holds that earlier events in a sequence are remembered more clearly [16,17]. Moreover, emotion can modulate these effects of primacy and recency [18]. Accordingly, the correlation between mystical experience and clinical efficacy is likely to be conditioned by the precise temporal dynamics and emotional content of the acute experience. There may be a vast repertoire of drug-induced experiential trajectories that are clinically useful, but remain undetected as we are unable to reliably measure them.

Such trajectories are difficult to characterize with traditional questionnaires. While questionnaires are indispensable, validated instruments for clinical research, they inevitably compress the vast range of psychoactive experiences to scalar values measuring only a few aspects of an experience (e.g. 14 for Hamilton Anxiety Scale [19] or 30 for the Mystical Experience Questionnaire [20] and CAPS-5 [21]). The complex dynamics of feelings throughout a drug experience are reduced to low-dimensional points in a space spanned by the researcher’s conceptual priorities, rather than those of the patient. In contrast, patient testimonials paired with large language models can rigorously quantify many dimensions of subjective experience as they evolve throughout a narrative report. These models unlock the possibility of data-driven neurofeedback protocols that customize psychoactive drug experiences and fine-tune them for desired outcomes. Here, we demonstrate 3 distinct modeling techniques (i.e. self-supervised, supervised and transfer learning) that all discover mutually-reinforcing and neurochemically-grounded structures from subjective reports.

Receptor affinities provide a neurobiological anchor for the phenomenology contained in the testimonials. A drug’s binding affinity for a receptor is not a proxy for its function; affinity values cannot distinguish between agonist and antagonist, or account for arrestin coupling, or biased intracellular signaling, or other factors that influence a drug’s effect. Nonetheless, the affinity data provides a common biochemical signature for each molecule in our study. Though the precise biological mechanisms of action cannot be inferred from these signatures, it is nonetheless possible to learn rich pharmacological representations from them. Likewise, the emotional landscapes richly depict retrospective associations, but are not necessarily predictive or mechanistic. Using these publicly-available data and large language models we show it is possible to build neurobiologically-informed, temporally-sensitive portraits of drug-induced subjective experiences. Such rich representations may be clinically useful even without knowing the precise neurobiological mechanisms underlying them.

Our contributions with respect to prior work that has explored Erowid testimonials using natural language processing [22–24] include: (1) extending [6] to cover 52 drugs and over 11,816 trips, eliminating affinities whose replicability had been questioned [25], while exclusively sourcing affinity data from the Psychoactive Drug Screening Program [26] thus registering a cacophony of pharmaceuticals into a unified, neurobiologically-grounded space of psychoactivity; (2) leveraging pretrained language models we demonstrate how both transfer-learning and supervised-learning enable modeling trajectories, rather than just static summaries, capable of discerning subtle similarities and differences between individual drugs and drug classes; (3) predicting demographics directly from natural language demonstrating the feasibility of automated bias reduction methods; and (4) demonstrating that these diverse modeling strategies all independently elucidate a common, striking structure that distinguishes the mystical lucid heights of psychedelia from the dysphoric, mundane, daily battles with addiction and mental illness.

## Results

We amassed a corpus of 11,816 psychoactive experiences from Erowid, which we semantically and chemically characterize with two BERT-based models, and one Canonical Correlation Analysis (CCA) see Fig. 1. The supervised model, BERTowid, is trained using multi-task, multi-label, classification and regression directly on Erowid testimonials and associated metadata. BERTowid is trained to “read” a 512 token excerpt from the testimonial and predict the associated drug, its chemical and pharmacological class, self-reported gender and age, 52 metadata tags, 11 canonical correlation component weightings, and 30 receptor affinities. Table 1 shows the taxonomy and testimonials counts. The transfer-learning model, BERTiment, is trained to detect 28 sentiments simultaneously on a corpus from Reddit [10]. It then makes inferences on Erowid revealing sentimental trajectories which we show agree with psychiatrist adjudications, validate expected emotional associations and are consistent with pharmacological groupings. Both models generalize to unseen data, demonstrating how machine learning on crowd-sourced, noisy semantic data can lead to diverse biochemical inferences. Note for instance in Fig. 2 how the entactogens MDA and MDMA, the opioids and the antidepressants all track together.

**Table 1:** *Drug taxonomy* The Drug taxonomy used in the study was sourced from the psychonaut wiki (https://psychonautwiki.org/).

| Drug | Chemical Class | Psychoactive Class | N Testimonials |
| --- | --- | --- | --- |
| 25i-nbome | Phenethylamine | Psychedelic | 150 |
| 2C-B | Phenethylamine | Psychedelic | 204 |
| 2C-C | Phenethylamine | Psychedelic | 64 |
| 2C-D | Phenethylamine | Psychedelic | 47 |
| 2C-E | Phenethylamine | Psychedelic | 303 |
| 2C-I | Phenethylamine | Psychedelic | 391 |
| 2C-P | Phenethylamine | Psychedelic | 58 |
| 2c-t-2 | Phenethylamine | Psychedelic | 118 |
| 2c-t-4 | Phenethylamine | Psychedelic | 16 |
| 2c-t-7 | Phenethylamine | Psychedelic | 173 |
| 5-meo-dmt | Tryptamine | Psychedelic | 281 |
| 5-meo-diPT | Tryptamine | Psychedelic | 208 |
| 5-meo-miPT | Tryptamine | Psychedelic | 71 |
| 5-meo-tmt | Tryptamine | Psychedelic | 109 |
| DMT | Tryptamine | Psychedelic | 530 |
| DOB | Amphetamine | Psychedelic | 45 |
| DOI | Amphetamine | Psychedelic | 34 |
| DOM | Amphetamine | Psychedelic | 36 |
| DPT | Tryptamine | Psychedelic | 148 |
| DXM | Morphinan | Dissociative | 457 |
| DIPT | Tryptamine | Psychedelic | 54 |
| Ketamine | Arylcyclohexylamine | Dissociative | 404 |
| LSD | Lysergamide | Psychedelic | 1177 |
| MDA | Amphetamine | Entactogen | 79 |
| MDMA | Phenethylamine | Entactogen | 1154 |
| Mescaline | Phenethylamine | Psychedelic | 136 |
| PCP | Arylcyclohexylamine | Dissociative | 76 |
| Psilocin | Tryptamine | Psychedelic | 597 |
| Salvia | Salvinorin | Hallucinogen | 199 |
| TMA-2 | Amphetamine | Psychedelic | 24 |
| Alprazolam | Benzodiazepine | Depressant | 120 |
| Amphetamine | Phenethylamine | Stimulant | 385 |
| Aripiprazole | Piperazine | Antipsychotic | 26 |
| Bupropion | Aminoketone | Antidepressant | 86 |
| Cocaine | Tropane_alkaloid | Stimulant | 568 |
| Diphenhydramine | Ethanolamine | Deliriant | 285 |
| Haloperidol | Butyrophenone | Antipsychotic | 12 |
| Hydrocodone | Morphinan | Opioid | 178 |
| Hydromorphone | Morphinan | Opioid | 46 |
| Ibogaine | Tryptamine | Psychedelic | 80 |
| Methadone | Diphenylpropylamine | Opioid | 110 |
| Methamphetamine | Amphetamine | Stimulant | 419 |
| Mirtazapine | Piperazinoazepine | Deliriant | 39 |
| Morphine | Morphinan | Opioid | 105 |
| Olanzapine | Thienobenzodiazepine | Antipsychotic | 46 |
| Oxycodone | Morphinan | Opioid | 237 |
| Paroxetine | Piperidine | Antidepressant | 77 |
| Quetiapine | Dibenzothiazepine | Antipsychotic | 146 |
| Risperidone | Benzisoxazole | Antipsychotic | 31 |
| Sertraline | SSRI | Antidepressant | 48 |
| THC | Cannabinoid | Hallucinogen | 1348 |
| Venlafaxine | SNRI | Antidepressant | 81 |

**Figure 1:**
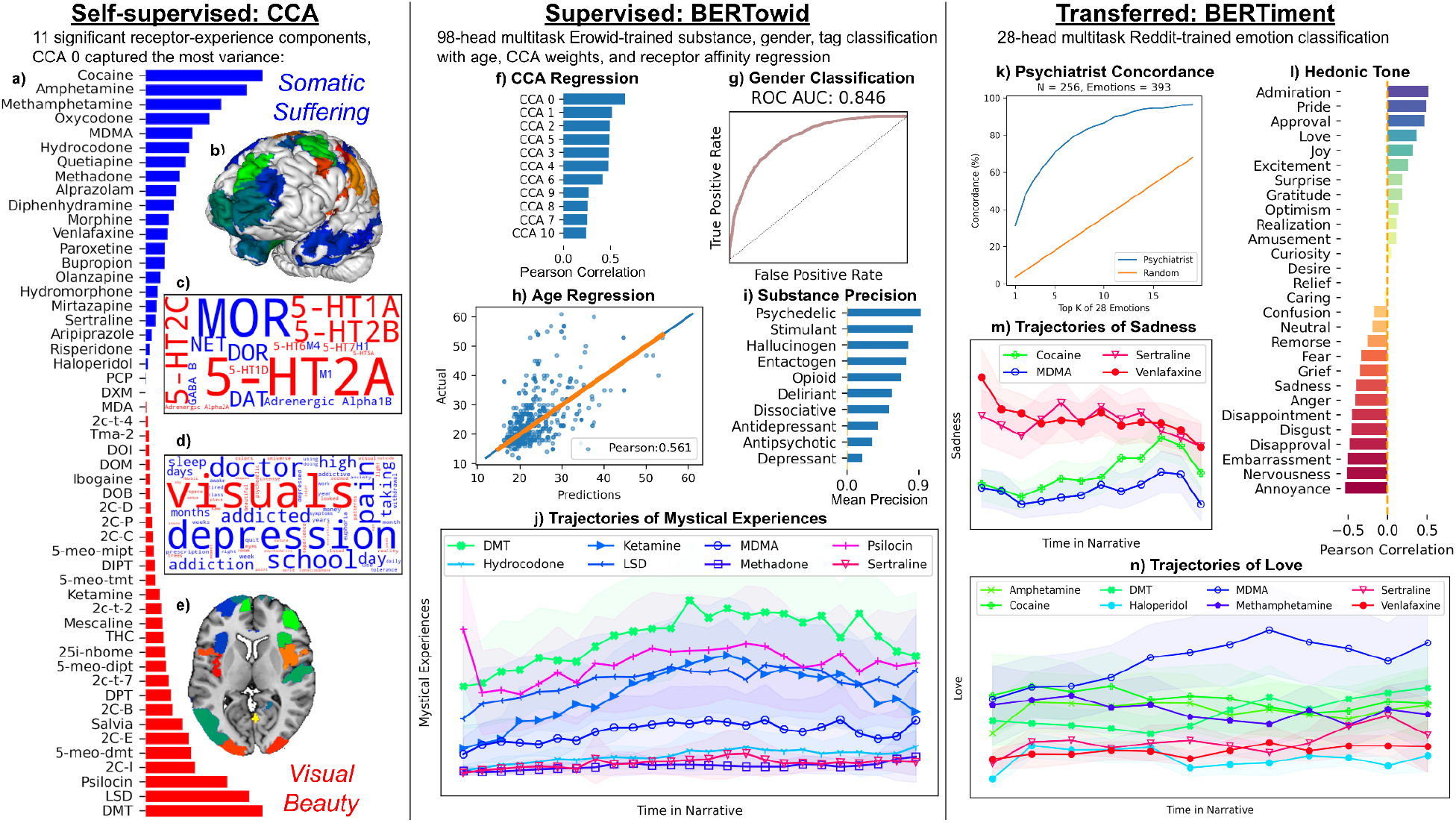
Models This figure illustrates the three models with a selection of key results.**Left:** the dominant component, CCA 0, found by the self-supervised learning CCA method. One extreme of CCA 0 encodes concepts of somatic suffering displayed in blue, while the other pole encompasses visual beauty and is displayed in red. All 52 drugs included in the study are shown in (a) with the ranking along CCA 0. (b) shows a brain surface map of CCA 0. (c) shows receptor clouds, note how the red visual/beauty pole is entirely serotonergic (5HT*) while the somatic/suffering pole in blue highlights several different neurotransmitter types including opioid (MOR, DOR), GABA and acetylcholine (M1, M4). (d) shows word clouds with font size determined by their CCA 0 weighting. (e) shows an axial slice of CCA 0 mapping into the brain, note how strongly the visual pole highlights the visual cortex. **Middle:** results from the supervised model BERTowid, which is a multi-task classification and regressing transformer trained directly on Erowid testimonials. All results are from test set testimonials, which were not used in training. (f) Pearson correlation with the 11 CCA factors per-testimonial weightings. (g) self-reported gender ROC curve. (h) Pearson correlation with self-reported age. (i) mean precision per psychoactive class. Tiling inferences from BERT models along the narrative of the testimonials we construct trajectories, for clarity we only show a few of 52 drugs here. (j) trajectories for the semantic tag of “Mystical Experiences”, note the prominence of DMT. **Right:** transfer-learning results from a model trained on an entirely different text corpus to classify emotions. (k) BERTiment’s concordance with a clinical-psychiatrist emotion adjudication in Erowid testimonials. (l) IMDB movie review hedonic-tone classifier correlation with the 28 emotions inferred on Erowid. (m) BERTiment Sadness trajectories, note how the antidepressants track with each other and are initially quite elevated. (n) BERTiment Love trajectories, note the prominence of MDMA.

**Figure 2:**
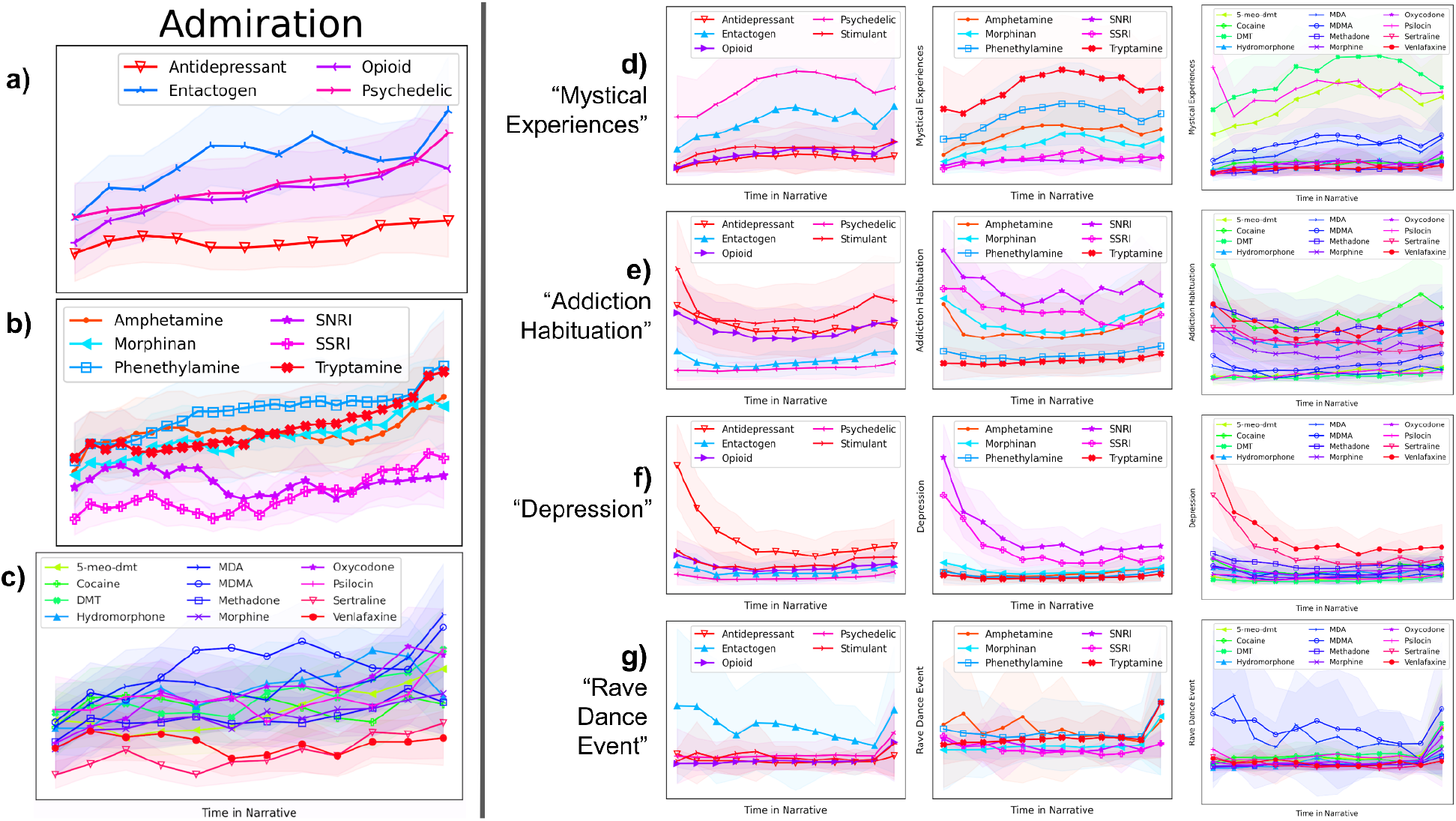
Trajectories and Drug Taxonomies **Left:** BERTiment trajectories for the emotion “Admiration” at the pharmacological level (a), chemical level (b) and individual drug level (c), see Table 1 for the full drug taxonomy. Note that for clarity we have selected only 12 representative drugs of the 52 included, see other figures for comparisons involving all drugs. **Right:** BERTowid trajectories for each of the 3 different levels of drug classification (from the left panel) on metadata tags “Mystical Experiences” (d), “Addiction Habituation” (e), “Depression” (f), and “Rave Dance Event” (g). Note the concordance between the entactogens MDA and MDMA and the antidepressants sertraline and venlafaxine.

This point is reinforced by our findings from CCA which identified a latent structure of 11 statistically-significant components mapping between the semantic data and the receptor affinity profiles in a self-supervised fashion [6]. CCA is a linear model and relies on a bag-of-words representation of the entirety of the testimonial text, while the transformers are deep nonlinear neural networks which positionally-encode a subset of text excerpted from the testimonials. Despite the large differences in representation and model, BERTowid learns to infer the CCA weightings, while many BERTiment emotion-scapes reveal similar drug rankings as given by CCA 0, see Fig. 3.

**Figure 3:**
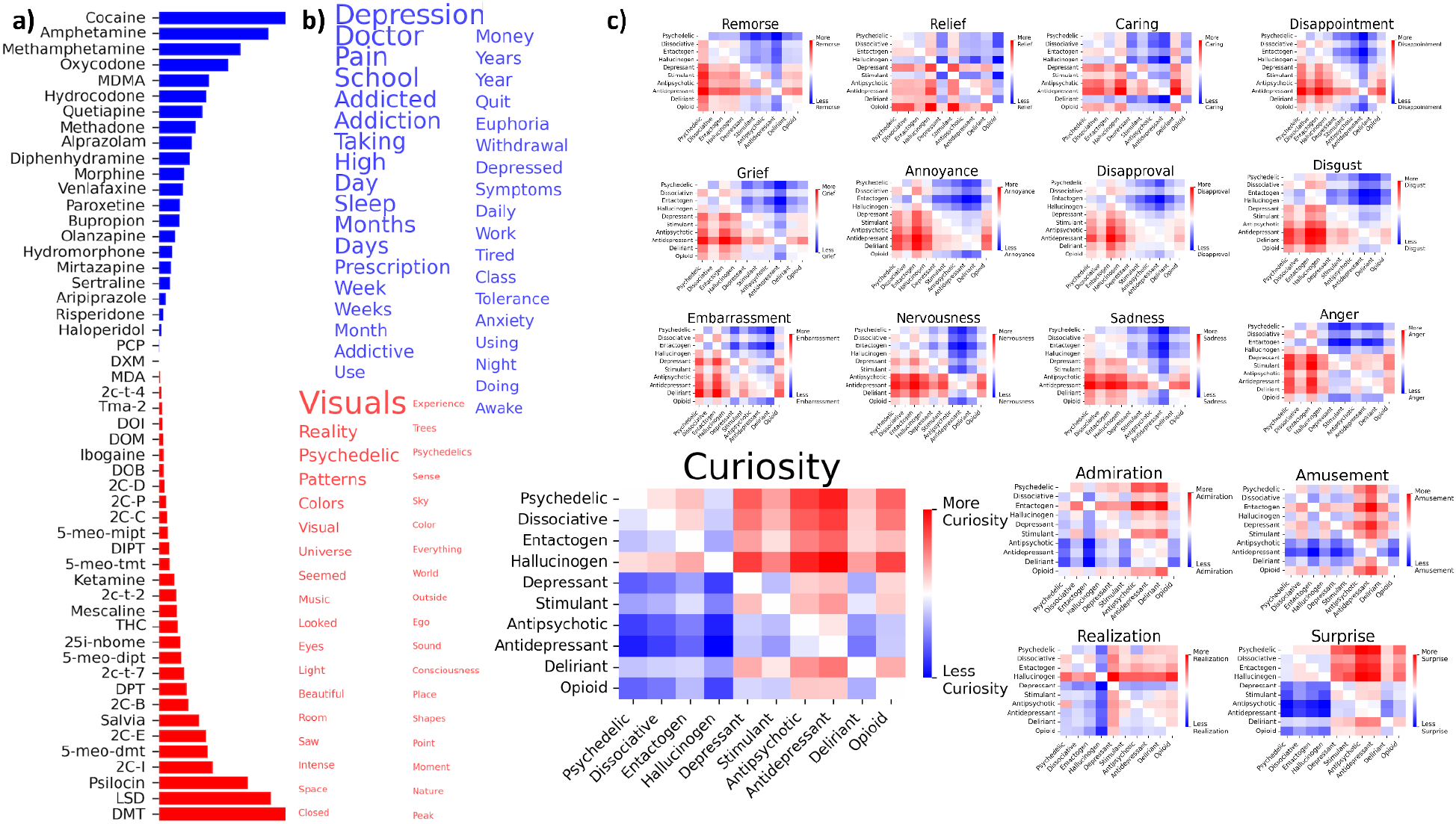
Dynamic Time Warp Distances Between 17 Emotions Supports CCA 0 CCA 0, the dominant component from CCA, revealed a ranking of the 52 drugs (a) whose semantic axis, shown by the words at each extreme in (b) were concordant with many emotion DTW matrices from BERTiment (c). Note how all 12 of the DTW matrices in the top half of the figure have bluish upper diagonals and reddish lower diagonals, while the opposite pattern was found for the 5 emotions in the lower half of the image.

### BERTowid

There is noise inherent in any crowd-sourced, open dataset like Erowid, which includes reports from many illegal substances rife with potential impurities and misrepresentations. Nonetheless, BERTowid shows powerful discrimination at several different granularities of pharmacology, classifying amongst 52 drugs, 22 ligand chemical types, and 10 pharmacologic classes, 30 receptor subtypes, and 11 CCA weights. Table 1 shows the taxonomies and testimonial counts. Model mistakes are consistent with expected biochemical and pharmacological groupings. For example, the psychedelic chemical classes of phenethylamines and tryptamines are much more likely to be mistaken for each other, than for a benzodiazepine, antidepressant, or opioid (see confusion matrices in Supplementary Fig. 1). Semantic tags are also learnable, with some of the best-performing being “Medical Use”, “Mystical Experiences”, “Alone” and “Addiction Habituation”, with areas under the receiver operating characteristic curves (ROC AUC) ranging from 0.88 to 0.95, areas under the precision-recall and ROC curves for all tags are shown in Supplementary Fig. 2.

Confirming its reputation as the “spirit molecule”, DMT displayed heightened trajectories for the tag “Mystical Experiences” and, even more dramatically, for the tag “Entities and Beings”, echoing themes uncovered in manual DMT-specific analyses [27]. As expected, the “Depression” tag trajectory highlights antidepressants, while the “Addiction Habituation” tag is consistently elevated for the stimulants cocaine and methamphetamine, see Supplementary Fig. 3.

Some testimonials include self-reported age (3,116) and gender (11,129) with which we trained gender-classifying (ROC AUC 0.85) and age-regressing (Pearson correlation 0.56) output heads. Despite missingness, gender class imbalance, and skew towards younger individuals, we can predict age and gender from these reports. Accurate detection of sensitive features, such as these, is a critical first step towards de-biasing predictions through iterative removal of confounded subspaces [28,29]. Given their potential role in healthcare, the ability to apply these models without bias is of utmost importance.

Through an entirely different analytic paradigm BERTowid appears to broadly confirm the salience and ranking of the 11 CCA components (described below). Test set performance on the CCA components drops off almost exactly with their ordering by CCA, with (Pearson correlations ranging from 0.68 for CCA 0 to 0.24 for CCA 11). Components which explain more variance between testimonials and affinity are also more effectively learned directly from testimonials.

### BERTiment

BERTiment is trained to predict 28 emotion classifications from Reddit annotations [10]. All emotion predictions generalize to unseen data with ROC AUCs ranging from 0.72 for “Realization” to 0.97 for “Love”. Further validation is provided by a hedonic tone classifying BERT model trained with positive and negative movie reviews from IMDB [12]. The signed Pearson correlations between BERTiment and the hedonic tone predictions neatly sorts the fine-grained emotion taxonomy. The emotions “Admiration”, “Pride”, “Approval”, and “Love” have the highest positive correlations, while “Annoyance”, “Nervousness”, “Embarrassment”, “Disapproval”, and “Disgust” have the largest negative correlations. Originating from entirely different datasets (i.e. movie reviews and Reddit posts) and evaluated on third orthogonal dataset (i.e. the Erowid testimonials) these models learned mutually reinforcing representations of sentiment, albeit at different levels of granularity, as shown in Fig. 1 panel (l).

Domain expert validation for the specific context of the emotions contained in reports of psychoactive experience was provided by a clinical psychiatrist, who manually adjudicated 393 emotions from 256 Erowid excerpts. Concordance between the model and the psychiatrist was within the range of inter-human variability as reported in the original GoEmotions paper [10]. Specifically, human labeled emotions were in the top 10 BERTiment emotions for 87% of the labels, in the top 5 for 73%, and in the top 1 for 42%, see Fig. 1 panel (k).

Qualitative manual inspection confirms that the extreme (positive and negative) predictions for each sentiment were prominent examples of the emotion (or its opposite), see Supplementary Table 1. As expected with extreme language, profanities, capitalization and modifiers like “very” and “so” are common. To quantitatively evaluate the sentimental trajectories, Dynamic Time Warping (DTW) measured the distance between the averaged trajectories for each emotion and each pharmacological and biochemical class, see Fig. 2 and Supplementary Fig. 4 [30]. The DTW reveals emotional landscapes that conform to expectations based on pharmacological classifications, molecular structure, questionnaires, and anecdotal reports of drug phenomenology [31,32]. The DTW matrices are skew-symmetric with the sign indicating which of the drugs had a higher mean predicted emotion. Fig. 4 provides a comprehensive view of the emotional content as determined by BERTiment in the Erowid dataset. Ordered by hedonic-tone, the most negative sentiments are associated with antidepressants and antipsychotics, in the middle we find pharmacological classes that are used clinically but also abused recreationally, like opioids and deliriants, and at the positive extreme we see psychedelics and entactogens. The association of psychiatric medications with negative emotions is confounded by ascertainment bias of those who seek out these medications, and does not necessarily reflect their efficacy.

**Figure 4:**
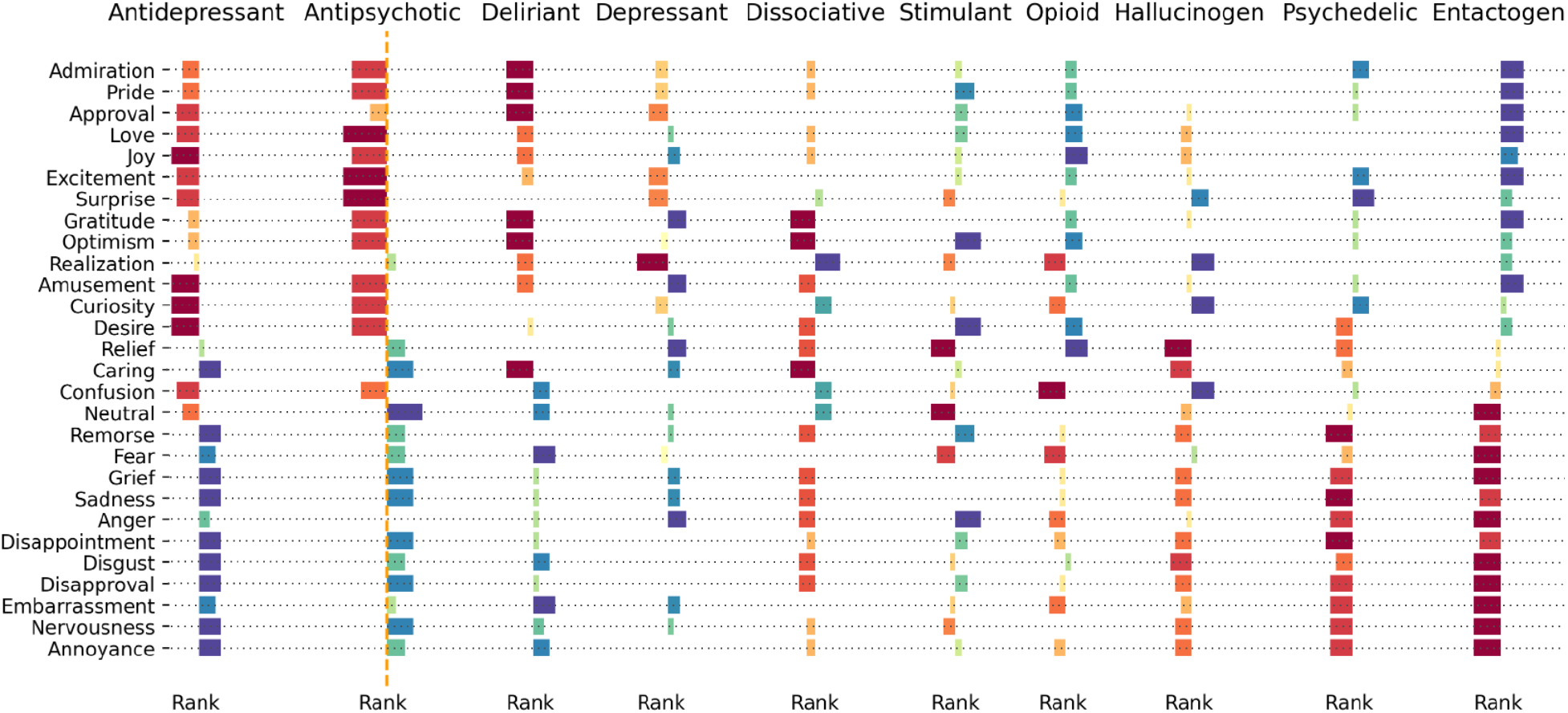
Emotionscapes The relative ranking amongst 10 pharmacological classes for BERTiment predictions for all 28 sentiments. The 28 sentiments are ordered by the hedonic tone spectrum derived from correlation with the IMDB movie review positivity classifier. Note that the drug classes are ordered from left to right according to the prevalence of negative vs positive overall hedonic tone, and that as expected antidepressants show the least hedonic tone and entactogens show the most.

Zooming in from the broad emotionscapes to a singular molecule, MDMA is characterized with both BERTowid and BERTiment in Fig. 5. The trajectory of “Love” during MDMA testimonials starts high and ends higher–fitting for a drug colloquially known as the “love-drug”. This arc is clearly distinguished from all other drugs, though closely tracked by the related entactogen, MDA. Supplementary Fig. 3 shows the “Sadness” trajectories of the stimulants (cocaine, amphetamine, and methamphetamine) are tightly coupled and rise gradually over the course of the testimonial, while the antidepressants (paroxetine, venlafaxine, and sertraline) start much higher than the stimulants but gradually fall. In contrast, the stimulants and antidepressants start with similar “Anger” levels, but over the course of the report methamphetamine and cocaine rise dramatically while for antidepressants they increase somewhat less.

**Figure 5:**
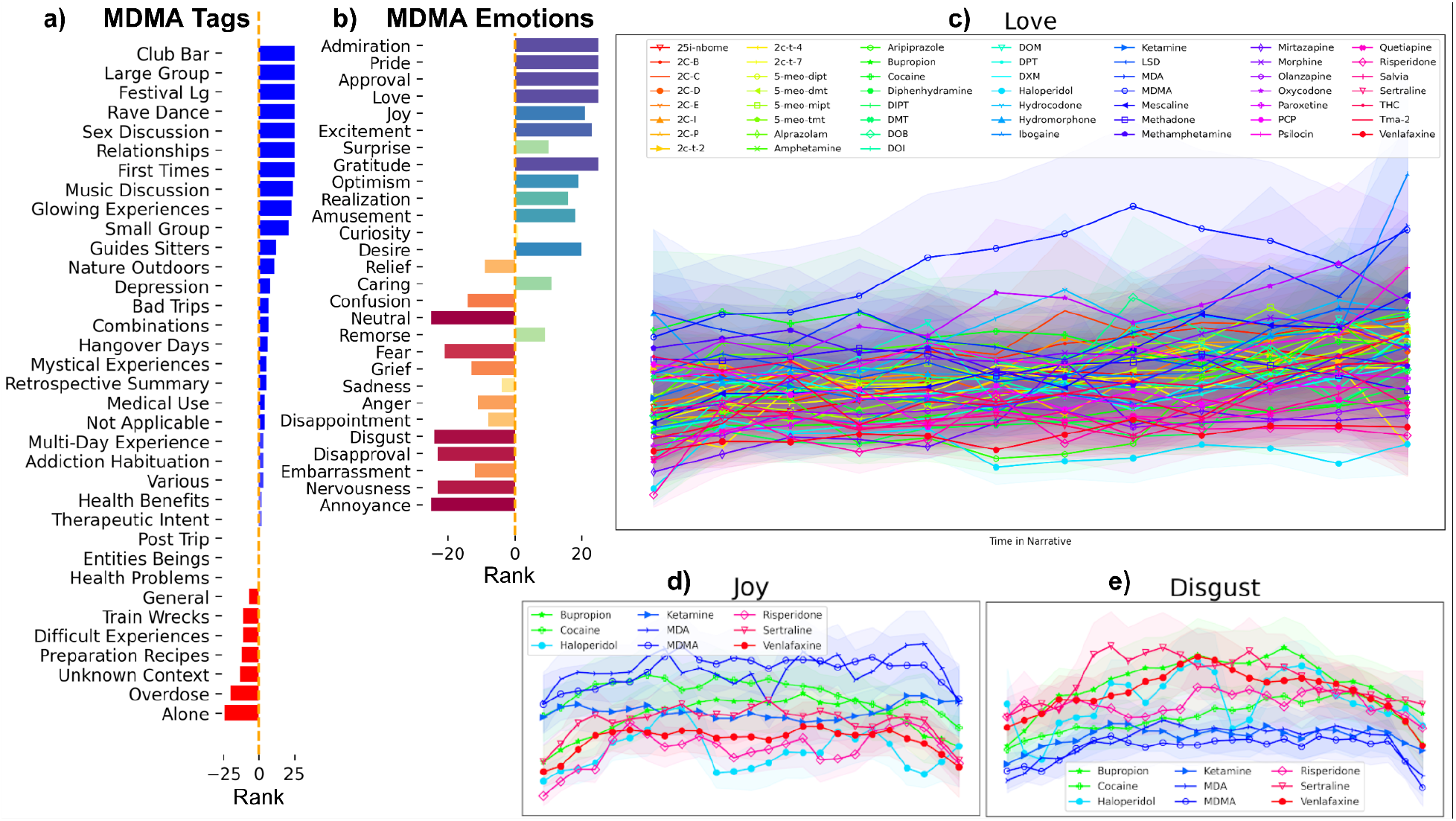
MDMA is Singular (a) The BERTowid tag predictions for MDMA compared to all other drugs, ordered from most associated (blue bars) to least associated (red bars). (b) BERTiment sentiment averages compared all other drugs, ordered by the hedonic tone spectrum, with higher values in blue/green, lower values red/yellow. (c)Trajectory of love compared against all drugs shows that MDMA is the dominant love drug. Sentiment trajectories for “Joy” (d), and “Disgust” (e) for MDMA along with representative drugs from the main pharmacologic classes are shown.

The emotions “Realization”, “Curiosity”, “Confusion”, “Surprise”, and “Amusement” are consistently elevated in subjective testimonials of hallucinogens and psychedelics as compared to other drug classes, most notably the opioids. This constellation of emotional trajectories provides additional discernment within the broad, overlapping classes of hallucinogens and psychedelics as shown in Fig. 6. For example, Salvia and DMT are both high in “Realization”, and “Curiosity”, however Salvia triggers more “Confusion”, while DMT generates more “Surprise”. PCP in contrast is high in “Confusion” and “Amusement”, but lower in “Realization”, “Surprise” and “Curiosity”. The opioids are consistently lower in all of these emotions, but “Relief” provides an interesting counterpoint, as it is higher in opioids than in hallucinogens or psychedelics, which one would expect for drugs widely prescribed for their pain-*relieving* effects.

**Figure 6:**
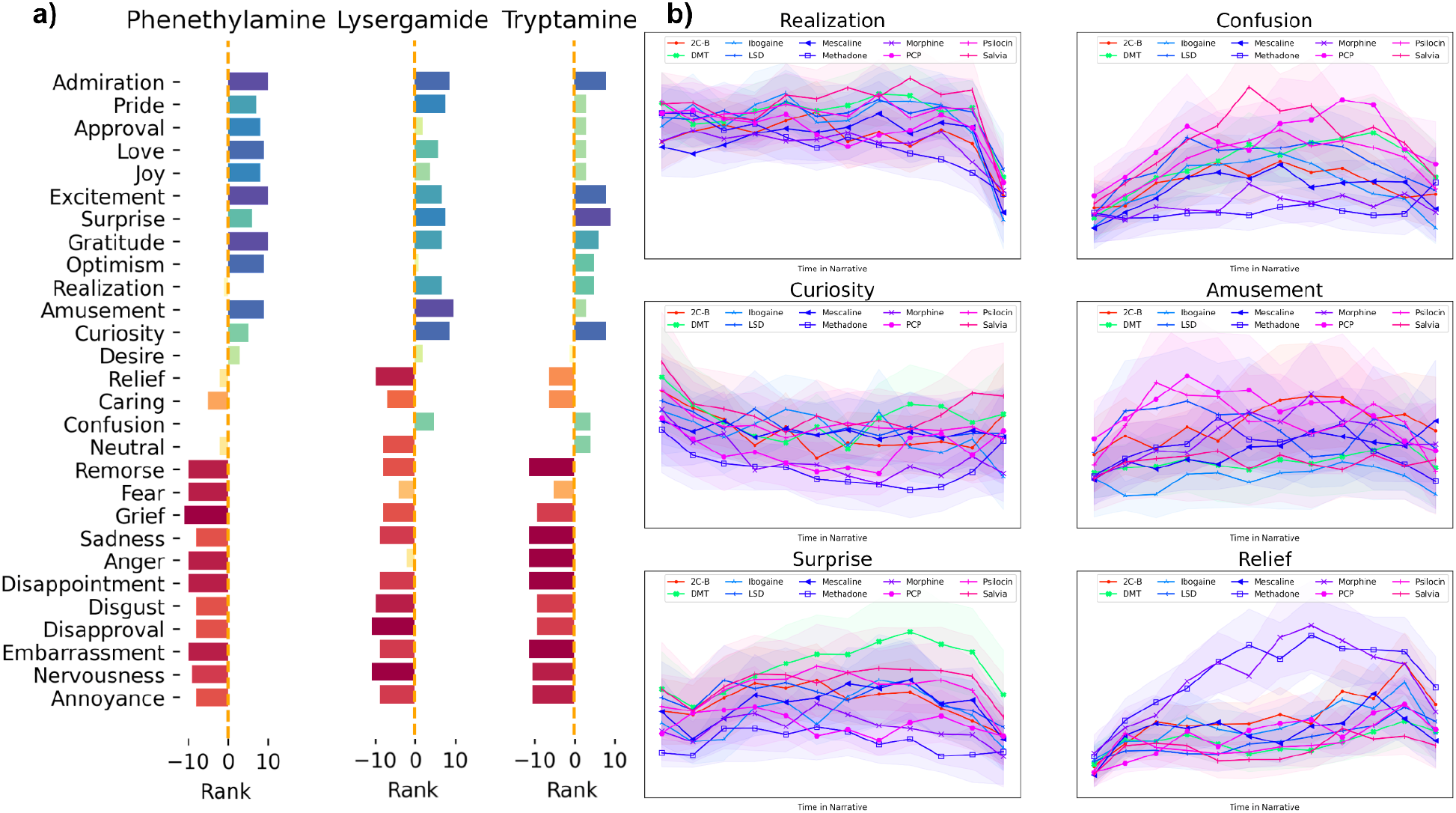
Psychedelics are Curious (a) the three major psychedelic chemical classes are shown with BERTiment’s relative ranking of each sentiment, as compared to the 22 classes included in the study. (b) BERTiment sentiment trajectories for a subset of phenethylamines, lysergamide, and tryptamines, and for comparison, opioids and ketamine are also shown. Note how, for tryptamines in particular, the weightings for “Surprise” and “Curiosity” are far greater than they are for MDMA. Whereas “Relief” is much higher for opioids than for any of the psychedelics.

Notably, not every emotion clearly distinguishes the drugs, the emotions “Neutral” and “Optimism” are quite conserved across pharmacological classes. For every drug analyzed, “Neutral” decreases as the testimonial proceeds, while “Optimism” trajectories for every drug increase dramatically near the end, resembling a ski-jump, see Supplementary Fig. 5. As if the peak-end rule is a self-fulfilling prophecy, testimonials for all drugs tend to end on an optimistic note. The reduction in “Neutral” over the course of a trip is expected as a drug’s effects reveal themselves to the user over time, a similar reduction in neutral sentiment over the course of a narrative was also shown with the IMDB [33].

### Canonical Correlation Analysis

Our pattern-learning strategy uncovered 11 discrete components that integrate subjective descriptions with the neurotransmitter affinity fingerprint of each psychoactive drug. The dominant factor described a vector characterized by *visual-beauty* on one extreme and *somatic-suffering* at the opposite. The second and third most explanatory factors contained poles of *depression-insomnia* and *impulsivity-addiction* contrasted with *perception-celebration* and *cosmic-expansion*, respectively. The eleven significant components reveal latent patterns of correlation between drug receptor affinity and word-usage, each is described in detail in Supplementary Figure: CCA 0 - Supplementary Figure: CCA 10.

Although not imposed by our analytic model, the cortical mappings of factors often turned out to be spatially contiguous with smooth transitions of expression strength between neighboring brain regions. These mappings were often found to be mirrored in homologous brain regions in the left and right hemispheres. These observations are noteworthy because all steps of our modeling pipeline were blind to brain regions and are informed only by receptor affinities and natural language.

The dominant component explained the largest portion of joint variation between receptor affinity profiles and word-usage frequencies. This central component isolated a semantic theme characterized by abstract, transcendental terms such as *reality, universe, everything, consciousness*, and *peak* as well as a constellation of perceptual phenomena including *visuals, patterns, music*, and *sound*. These terms were linked to drugs DMT, LSD, and psilocin, as well as mostly serotonergic receptor affinity at 5-HT2A, 5-HT1A, 5-HT2C, and the expression of these receptors in the medial prefrontal cortex. The opposite extreme of this factor flagged words that describe a theme of mundane suffering: *depression, pain, addiction, sleep, awake, daily, weeks, months, work, and anxiety*. This latter theme was linked to drugs cocaine, amphetamine, methamphetamine, and oxycodone, as well as receptor affinity at MOR, DOR, NET, and DAT, while the expression of these receptor genes was densest in the posterior cingulate cortex and the inferior parietal lobule. Notably, this component evoked a similar structure found by the BERTiment model for 17 of its 28 emotions, see Fig. 3.

### Convergences Between BERTiment, BERTowid, and CCA Components

Remarkably, the three distinct ML approaches elucidated similar findings. MDMA was found to elicit a uniquely positive palette of affective attributes that are of particular interest given its apparent efficacy as a treatment for PTSD [34]. At the most positive extreme of the IMDB-derived spectrum of hedonic tone, MDMA was ranked highest among all 52 compounds for “Admiration”, “Pride”, “Approval”, “Love”, “Excitement”, and “Gratitude”, and was only narrowly edged out by opioids for “Joy”. BERTowid tag weightings for entactogens (Supplementary Figure 11 and 12) exhibited heightened levels of festivity and dancing, but also relational, emotional, and effusive, (e.g. “Glowing”) phenomena. Using CCA to incorporate its receptor-affinity fingerprint, MDMA was most closely linked with receptor-experience patterns highlighting *perception*-*celebration* and *emotional-extremes*, with receptor gene-expression in the visual and primary sensory cortices.

Results also show important distinctions amongst psychedelic subclasses. In particular, the tryptamines exhibit elevated levels of “Curiosity”, “Surprise”, and “Realization”, while phenethylamines highlight relatively higher levels of “Admiration”, “Excitement”, and “Gratitude”. Tryptamines–especially powerful, short-acting DMT and 5-MeO-DMT–had higher “Mystical Experiences” tag weightings than the phenethylamines like mescaline and 2-CE. Interestingly, the chemically distinct diterpenoid compound Salvinorin A shared with these prototypical tryptamines high levels of “Surprise”, “Curiosity”, and “Realization”, in addition to a very high ranking of “Mystical Experience”.

Drug factorization in the dominant CCA component (CCA 0) aggregated stimulants (including MDMA) and psychiatric medications into one pole, and associated them with phenomena relating to suffering, addiction, and the mundane. Through an entirely separate analysis, BERTiment DTW charts demonstrated these drug classes all ranked highest for a constellation of emotions compatible with this CCA theme such as “Disappointment”, “Grief”, “Annoyance”, “Disapproval”, “Disgust”, “Sadness”, and “Anger”. The opposite pole grouped psychedelic and hallucinogenic drugs together with terms highlighting a theme of abstract, lucid, and beatific phenomena. BERTiment DTW found these drugs to have much higher scores for “Curiosity”, “Admiration”, “Amusement”, “Surprise”, and “Realization”. This dichotomy between the gloomy, prosaic, quotidian aspects of human sentiment and the more abstract, expansive, and creative aspects of human potential was judged by the CCA to be the most explanatory distinction in this large corpus of drugs, and the majority of sentiment dimensions fit neatly into the same schema.

## Discussion

Utilizing linear CCA and nonlinear transformers, supervised or self-supervised, we’ve trained models which represent diverse drug experiences in a unified, biochemically-informed, temporally-sensitive way. Derived directly from natural language, these representations contain information like the intensity of mystical experience and the depth of joy, anger, and grief: qualities of subjective experience that are of great clinical import for psychiatry. Sequentially applying these models on retrospective reports creates evocative narrative trajectories, which reflect pharmacological distinctions and conform to expectations of subjective effects reported by psychonauts and researchers. Our findings also generally dovetail with prior efforts to use natural language processing tools to analyze the Erowid testimonial database. For example, one recent study[35] found, in agreement with our results, that antidepressants were associated with words denoting negative affect and less with mystical phenomena–which we both attributed at least in part to ascertainment bias of depressed subjects seeking medication. Also, a pioneering study from 2012[22] noted surprising similarity between pharmacologically distinct but similarly short-acting drugs like Salvinorin A and DMT. Our analysis detected strong similarities between these drugs, but we were further able to parse fine-grained subjective differences with DMT being linked to relatively higher levels of “Surprise” while Salvinorin was associated with more “Confusion”.

Notably, compared to the aforementioned prior efforts, the ability to construct detailed, trajectories of experience using transformers that we introduce in this paper is a novel addition to the field. The peak-end rule inspired us to look for such trajectories since it established the counterintuitive finding that there are times when more pain is preferred to less. But this is a rule not a law, and exceptions such as whether pleasure is increasing or decreasing when an experience ends can be even more important than the peak level of pleasure felt [36]. There may well be further variations, and subtleties by which experiential trajectories shape how memories are formed. The shapes of the trajectories produced by BERTowid and BERTiment capture this nuance and, if combined with clinical outcome data in a prospective manner, may identify more temporally-mediated “rules” that mediate how drug-induced experiences impact overall mental health.

While the trajectories we constructed unfold from the narrative language in the trip report, the methods we describe naturally apply to other streams of phenomenological and neurochemical data. Modalities as diverse as EEG, ECG, fMRI, and other biometrics are amenable to trajectory construction, simply by replacing the BERT models with appropriately pre-trained encoders (e.g. a 1D CNN for EEGs or ECGs [37,38]). Excitingly, such modalities can be sampled with high temporal resolution and independently from the patient’s recollection, mitigating the issues inherent in self-reported datasets like Erowid, where there is uncertainty about dosage, chronology, and drugs’ impact on memory [39,40]. On the other hand, given that recent machine learning models trained on cross-modal representations have shown improved phenotype prediction [29], combining parallel subjective and neuroimaging datastreams may build more useful and holistic representations of acute drug states. By tethering retrospective linguistic reconstructions of drug experiences to specific timepoints in their corresponding EEG trajectories we can construct powerful cross-modal representations of conscious states.

Such cross-modal representations that provide a common measure spanning psychiatric illness, pharmacological intervention, and acute psychedelic experiences could be transformative to clinical practice. Combining real-time datastreams with the zero-shot learning capability of transformers [13,41], it is possible to envision a future of personalized, responsive psychoactive sessions. Music appreciation has been shown to be enhanced by certain psychedelics [42], but what if therapists could monitor and adjust the emotional palette as it is felt in real-time? The tables of extreme predictions from our models (Supplementary Tables 1 & 2) make clear that transformers learn semantics reflecting both the training label as well as its opposite meaning.

Word2Vec demonstrated that directions, not just points, are meaningfully encoded in latent spaces of natural language; (e.g. King-Queen = Man-Woman) [43]. The directions connecting the extremes of “Love” or “Mystical Experience” can guide trips as they unfold, with realtime neurofeedback modulating diverse environmental inputs. In this manner, psychedelic therapists of the future may use machine learning to help their patients navigate latent spaces of subjective experience towards psychoactive journeys of minimal risk and maximal therapeutic benefit.

## Online Methods

Many datasets were leveraged in this study, namely the Erowid dataset of 11,816 drug testimonials [44], receptor affinities at 61 receptor subtypes for 44 drugs from the Psychoactive Drug Screening Program (PDSP) [26,45] (Supplementary Figure 10), augmented with affinities for 8 phenethylamines from Rickli et al. [46], RNA gene expression data for 200 brain regions from the Allen Brain Atlas [47], and 58K Reddit posts with 28 human-annotated human emotions [10]. The 10 pharmacologic classes and the 22 chemical classes were retrieved from the Psychonaut Wiki [48].

### Data Preprocessing for Transformers

Scraped testimonials were parsed for meta data, drug-masked, and tokenized. Drug-masking removed all occurrences of drug names in the testimonial text, including both scientific, common, and colloquial nomenclature as well as misspellings. See Supplementary Table 3 and code for the full list of masked words. Models were initialized with pre-trained weights for the base BERT encoders. All initial model weights are publicly available. Except when otherwise noted, the base BERT model used was trained with the Stanford Sentiment Treebank data [33] and is available at TensorFlow Hub. The pooled output from the base BERT model was extended with a dropout layer [49] followed by a dense layer for each task (e.g. BERTiment has 28 distinct outputs–one for each binary emotion classification: present or absent). Code necessary to replicate our findings is available at: https://github.com/lucidtronix/bertowid

While testimonials vary in size, the input to BERT models is at most 512 tokens. A sliding window inference step used all available data by creating prediction series of varying lengths for each testimonial from each model. Different window sizes are compared in Supplementary Fig. 6. When the window size exceeds the testimonial size, the input is zero-padded. When the window size is smaller than the testimonial size, the testimonial is split into contiguous blocks of text and the model is applied to each on constructing a trajectory of inferences. Dynamic Time Warping (DTW) quantified inter-trajectory distances, using an implementation from the fastdtw python package [50]. The Broad Institute’s ML4H tools were used for model evaluation [51].

### BERT-based model encoder fine-tuning

The encoder backbone of both BERTowid and BERTiment is a bidirectional transformer (BERT) trained with a masked language model objective [52], [14]. BERTowid and BERTiment add output heads, which take the BERT encoder representation as input and fine-tune it for new tasks. The encoder backbone contains 109 million parameters. Base models are compared in Supplementary Fig. 7, showing similar performance. Prior to the output layer for the fine-tuning task, we insert a dropout layer, see Supplementary Fig. 8 [49]. The ADAMw [53] stochastic gradient descent optimization with initial learning rate of 1e-5 and a batch size of 32. The minimum validation loss model is serialized for downstream inference after 16 epochs.

### Training BERTowid

The Erowid metadata, the inferred CCA weights and the receptor affinities from PDSP provide diverse training labels for BERTowid. Both classification (e.g. drug, tag, gender) and regression tasks (e.g. age, affinity, CCA weights) are considered. All classification tasks are trained to minimize a cross entropy loss, while regression models are trained to minimize the mean squared error of their predictions. Classification and regression with a single model requires a term to balance between the two types of loss, but optimization was found to be sensitive to this value, requiring careful tuning to avoid convergence for only one of the loss types. To mitigate this, multitask BERTowid is trained and serialized separately for classification and regression. Supplementary Fig. 16 compares multi-task vs single task models showing a relatively small cost to taking the multitask approach. With a window size of 64 words and all categorical tasks, 16 epochs takes about 3 hours on a NVidia V-100 GPU.

Many of the categorical labels are class-imbalanced. This imbalance leads to poor performance on the less well-represented drugs and tags. To mitigate this we considered a weighted cross entropy loss, which scaled the loss by the inverse of each labels’ prevalence to compensate for the imbalance. As Supplementary Fig. 17 shows, the weighted loss did increase precision for less well represented drugs (at the cost of reduced precision on the more prevalent drugs). However, the results for tags were less convincing with only minor improvements in precision for the 4 rarest tags. A likely explanation is that less common tags are in fact less informative and may be applied less rigorously or consistently by the Erowid moderators. Giving these less informative labels more weight in the loss function results in a worse model. With a window size of 64 words and all categorical tasks, 16 epochs takes about 3 hours on a NVidia V-100 GPU.

### Training BERTiment

The BERTiment training procedure has previously been described and evaluated [10]. Our approach only differs in dropout rate, 0.5, learning rate, 1e-5, and batch size, 32, where for consistency we used the same hyper-parameters and base model used to train BERTowid. The GoEmotions data contains about 58K Reddit posts and 28 emotion annotations, which can be downloaded here. We split the data 70-20-10 between training, testing and validation sets. The 28-head BERTiment is trained for 16 epochs which takes about 2 hours on a NVidia V-100 GPU.

### Canonical Correlation Analysis

The CCA mapping between testimonials and affinities has previously been described in detail [6]. The approach here extends from 27 drugs to 52, from 40 receptor subtypes to 61, and exclusively sources affinity values from the PDSP [45] or Rickli et al. [46]. Each of these receptor-semantic components were composed of two poles of a weighted list of words and a weighted list of receptors. The receptor weights for each component were mapped to the cortex using Allen Brain Atlas receptor RNA expression quantities measured by invasive tissue probes. The scikit-learn package’s implementation of the CCA algorithm was used [54].

## Acknowledgments

We thank the founders, curators, contributors, and volunteers of Erowid Center for sharing data and for decades of work on the experience report collection.

## Author Contributions

S.F.F and G.B. conceived of the study and wrote the manuscript. S.F.F. wrote the code to train and evaluate the models, conducted the statistical analyses and created the figures. G.B. annotated the emotions from testimonial excerpts.

