## Supplemental Material for "Language Models Learn Sentiment and Substance from 11,000 Psychoactive Experiences"

### Supplemental Information for Language Models Learn Sentiment and Substance from 11,000 Psychoactive Experiences

#### Supplementary Figure 1: Drug Classification Performance

Test-set areas under the precision-recall and and ROC curves as well as confusion matrices for pharmacological class, chemical class, and individual drug labels are shown here.

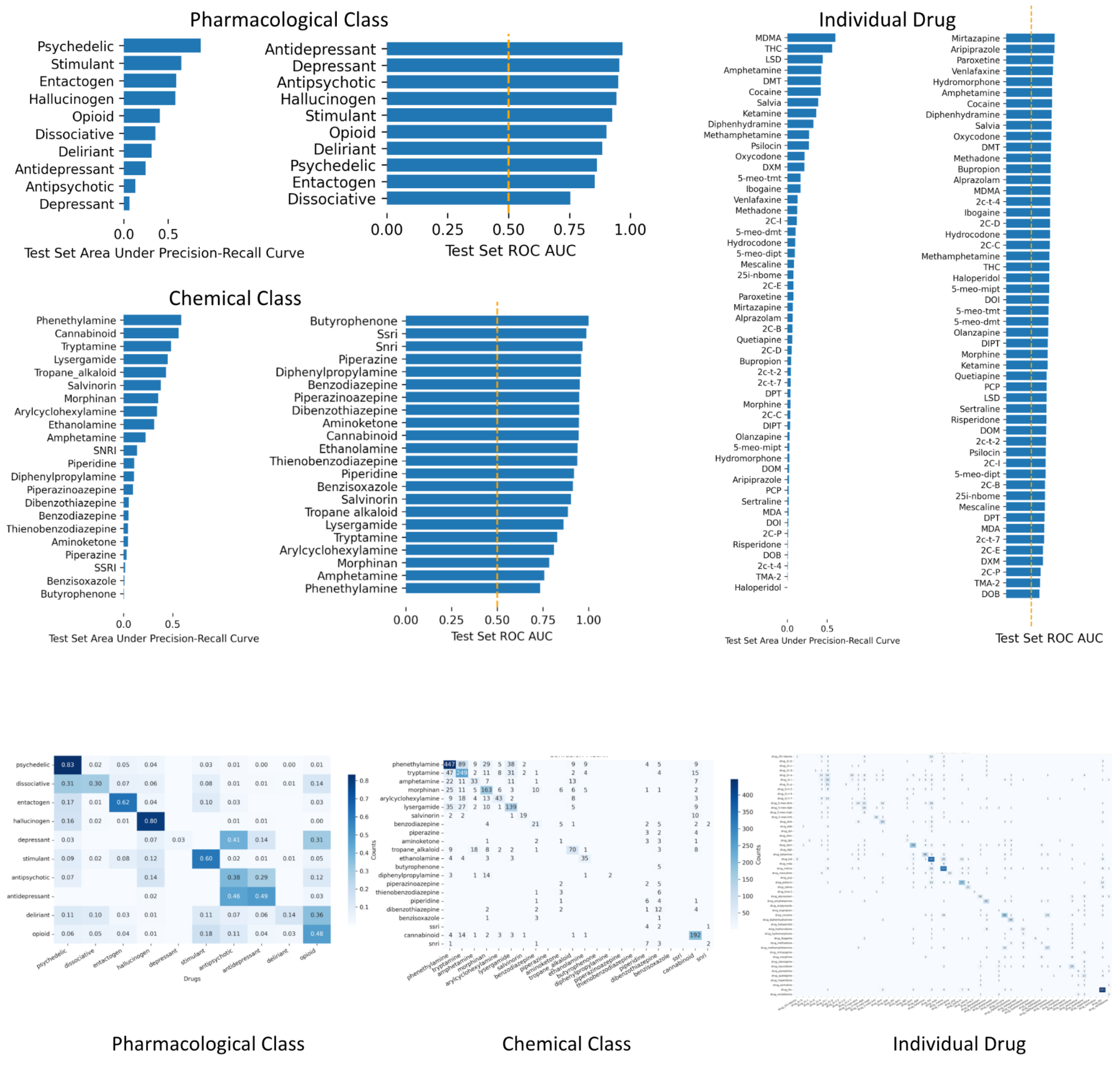

### Supplementary Figure 2: Results

BERTowid multitask evaluation on held-out test data and BERTiment comparison with clinical psychiatrist emotion adjudications and generalization to hedonic tone on Erowid testimonials. BERTowid classifies drugs at 3 different levels, amongst the 10 pharmacologic classes (A), 22 chemical classes (B), and the 52 drugs included in the study (C). The ROC AUC for the 35 meta tags with positive examples > 150 are shown in the third column (D). Test set Pearson correlation of the 11 CCA weight predictions (E) and the 30 receptor affinities (F). The ROC curve for self-reported gender (G), and the scatterplot for self-reported age (H) are shown in the bottom left. BERTiment evaluations are in the rightmost column with comparison with psychiatrist adjudicated emotions top right (I), and Pearson correlation with hedonic tone prediction on the Erowid dataset from the IMDB trained positive sentiment BERT classifier (J) at bottom right.

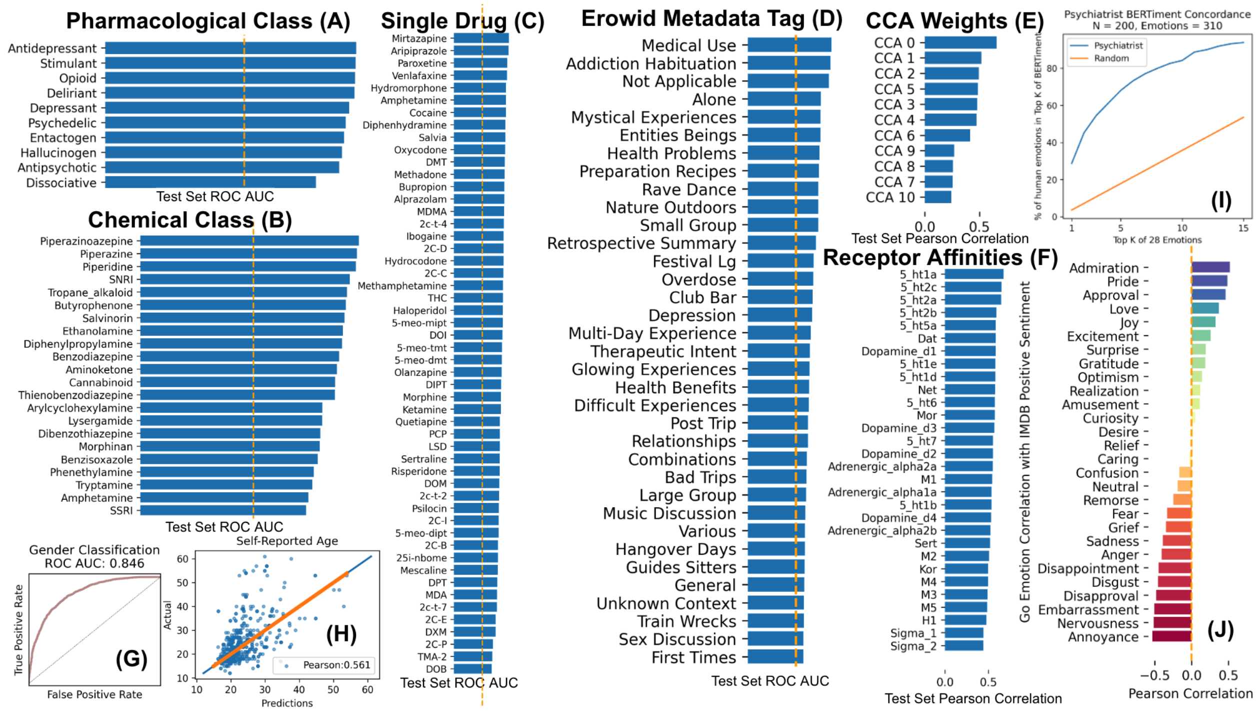

### Supplementary Figure 3: Trajectories

Applying BERTowid (left plots) and BERTiment (right plots) in a sliding window across each testimonial followed by linear interpolation gives the trajectories shown here. The first row shows the “Addiction Habituation” trajectory from BERTowid and the “Anger” trajectory from BERTiment, note how the stimulants cocaine and methamphetamine correlate with each other and between the models. This convergence is even clearer in the second row with BERTowid’s “Depression” tag and BERTiment’s “Sadness” label.  Antidepressant testimonials start high in both and trend down, while stimulants start much lower but trend upward.  Rows 3 and 4 show how the Erowid tags of “Mystical Experience” and “Entities Beings” flag the “spirit” molecule, DMT, while the emotions “Love” and “Approval” highlight the “love” drug, MDMA, as well as the related entactogen, MDA.

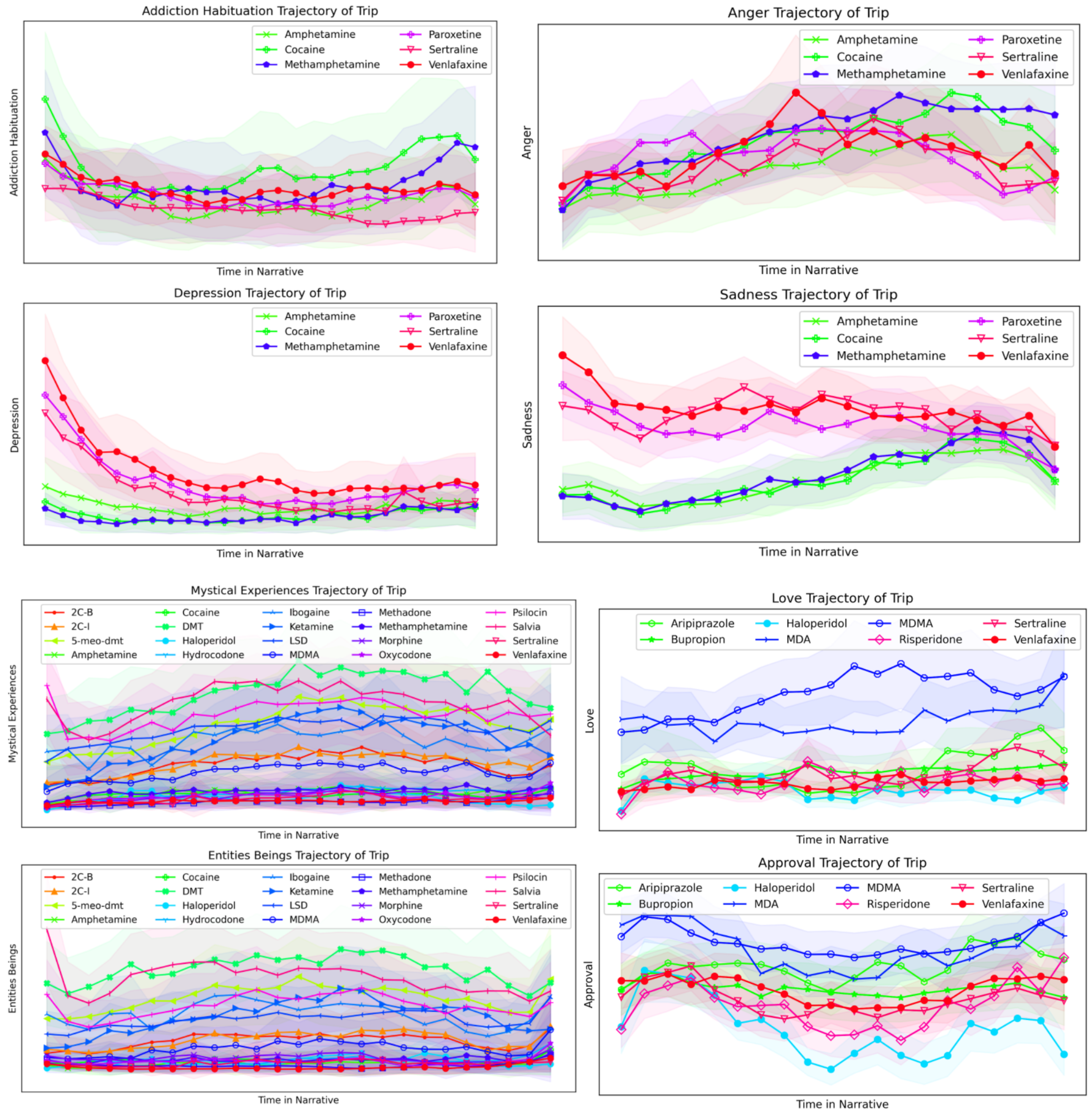

### Supplementary Figure 4: Compare Trajectories with Dynamic Time Warps

To achieve a comprehensive view of the emotional trajectories we create cross-drug DTW distance matrices where each cell contains the DTW distance between the drug (or drug class) in the row and in the column.  We use a signed distance which is positive (shown in red) if the mean sentiment prediction of the row's drug is higher than that of column's drug and negative (shown in blue) otherwise.  BERTiment sentiment trajectory for "Love" is shown for all drugs. MDMA clearly has the highest value through the majority of the narrative of drug experience. The chart on the upper right confirms this through DTW with MDMA creating a bright red line of "Love" slicing horizontally across the center. DTW by pharmacologic class shows entactogens (MDMA & MDA), compared to all other pharmacologic classes, have relatively low values for "Grief", high values for "Joy" and "Love", and moderate/high values for "Curiosity". DTW for BERTowid tag trajectories found several categories dominated singularly by MDMA ("Festival-Crowd", "Large-Group", "Sex-Discussions", "Rave-Dance", "Club-Bar").

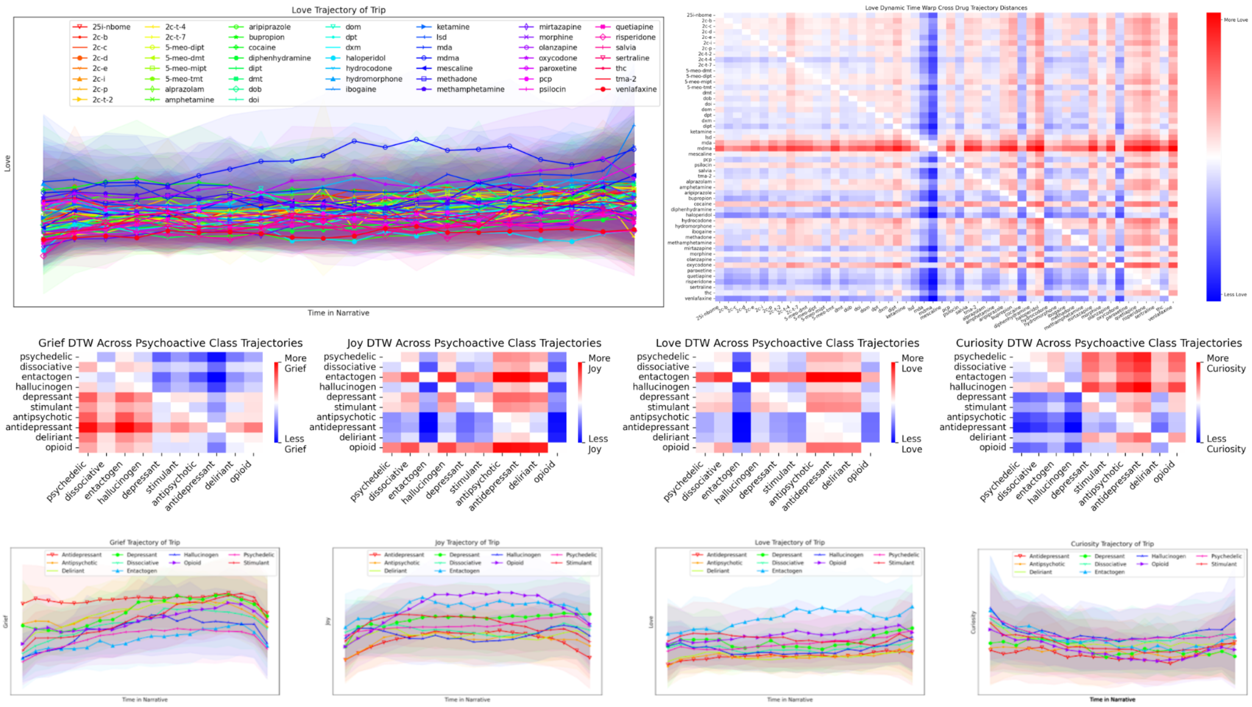

### Supplementary Figure 5: Optimism and Neutral Trajectories

While many sentiment trajectories showed substantial differences between drugs and drug classes, trajectories from the sentiments “Neutral” and “Optimism” were remarkably uniform across all drug groupings. “Neutral” gradually decreases over the course of the narrative. In contrast, “Optimism” showed an early, uniform rise then stayed steady for the most of the drug experience before uniformly increasing at the end.

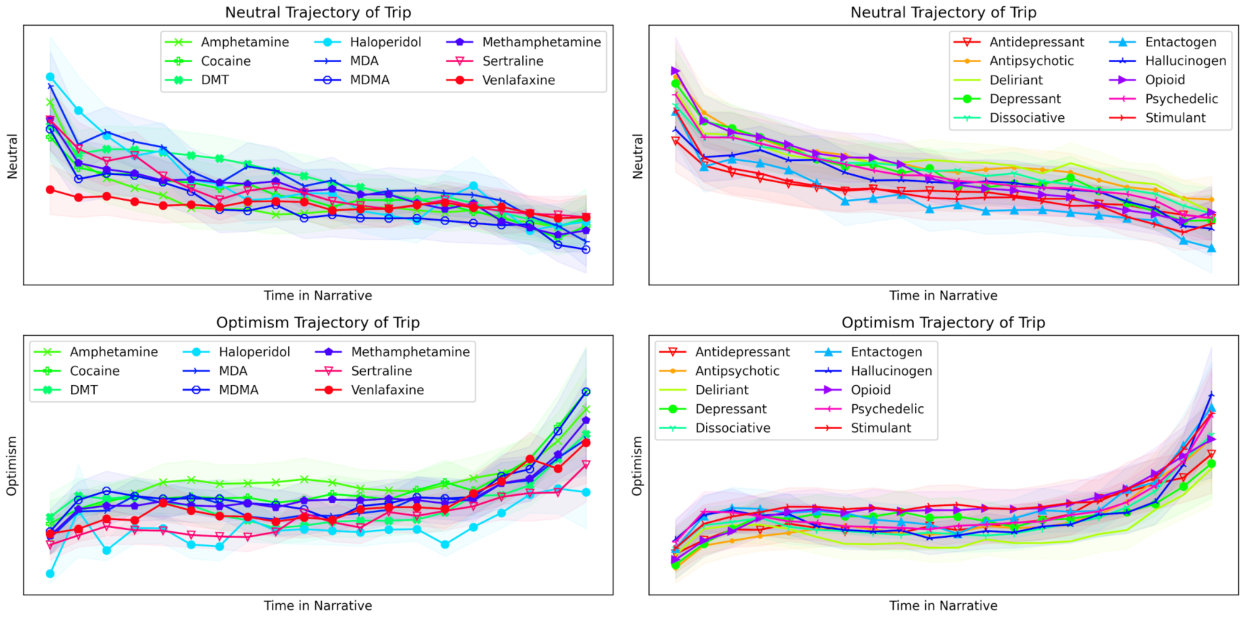

### Supplementary Figure 6: Word Window Size Comparison

Comparison of word size maximums for different BERTowid models on drug and metadata tag classification.  Maximum window sizes of 64 words were used for the results in the main paper.

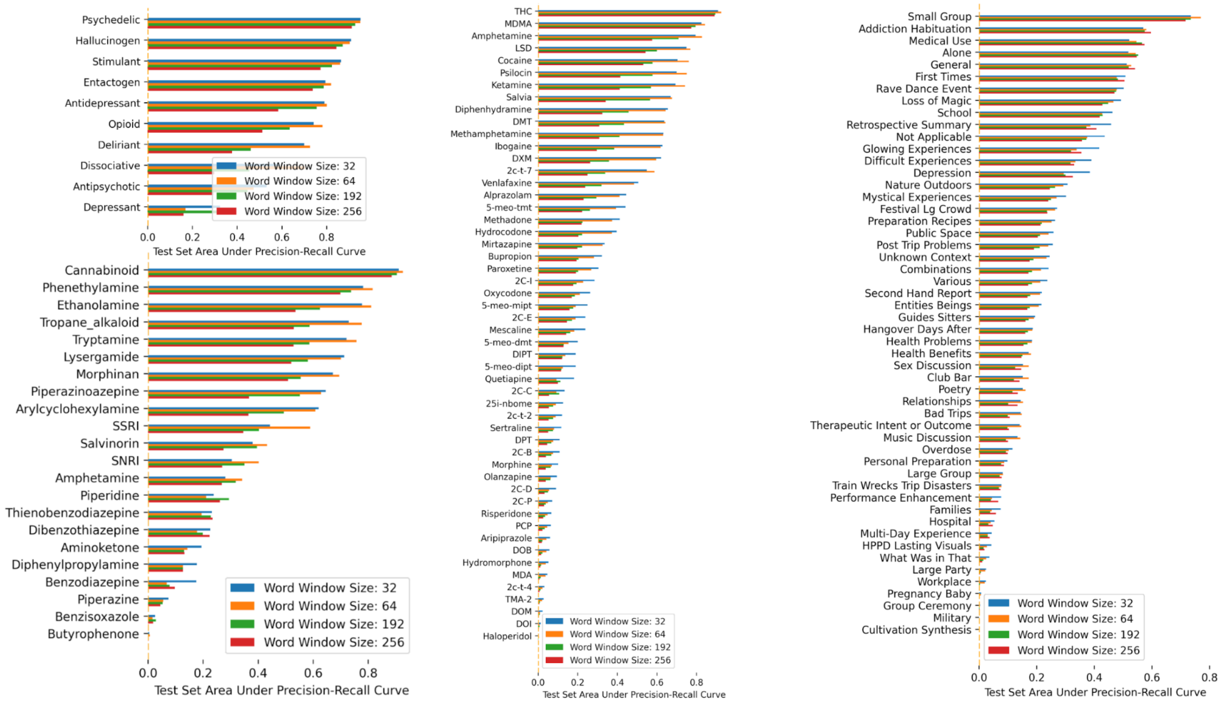

### Supplementary Figure 7: BERT Encoder Weights Comparison

To ensure optimal performance, standard BERT encoder weights were compared weights from SST2 and comparable AUC for Precision-Recall was comparable, and SST2 weights were ultimately used for the entire BERTowid analysis.

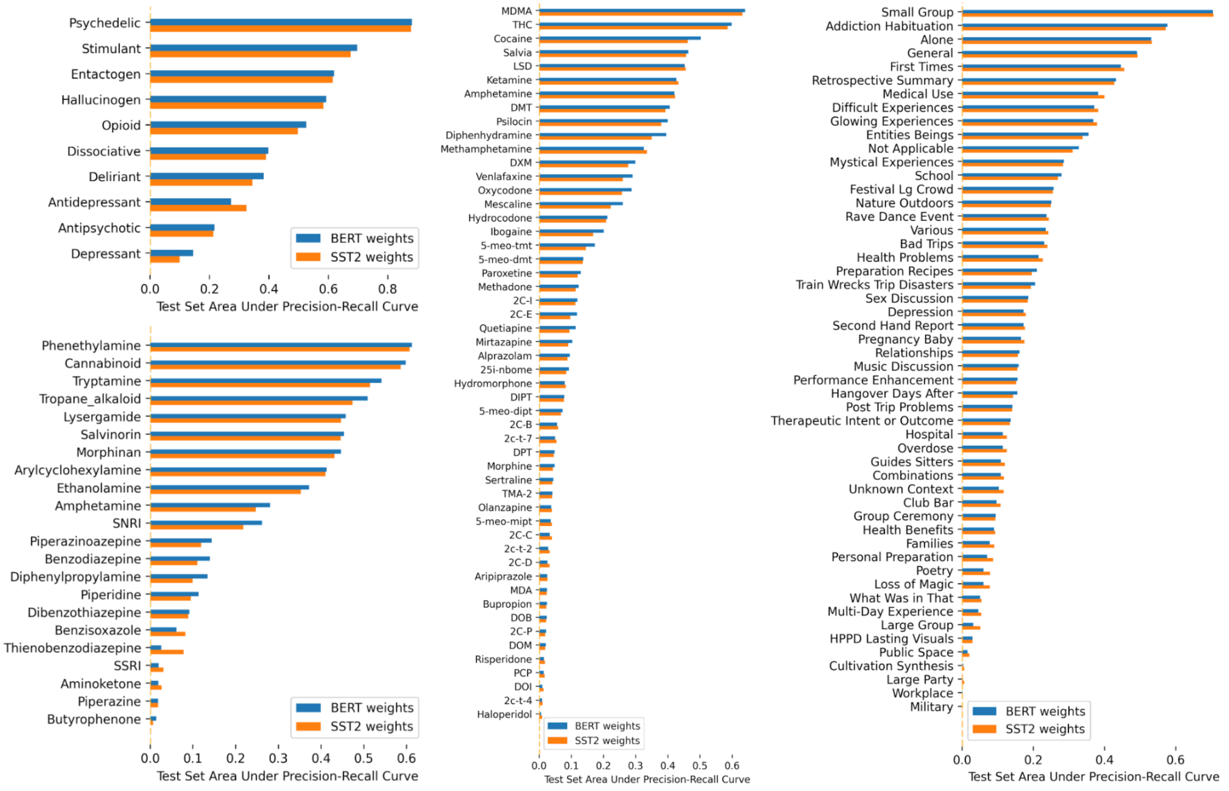

### Supplementary Figure 8: Dropout Rate Comparison

Dropout rates of 0.2, 0,5, and 0.7 perform similarly with best generalization results at 0.5. The learning curve comparison at far right shows the multitask loss at the end of each training epoch for the training set in blue, and the validation set in orange.  Notice how increasing the dropout rate delays overfitting.  The test set area under the precision recall curves are computed from a separate hold out set distinct from the training and validation testimonials.

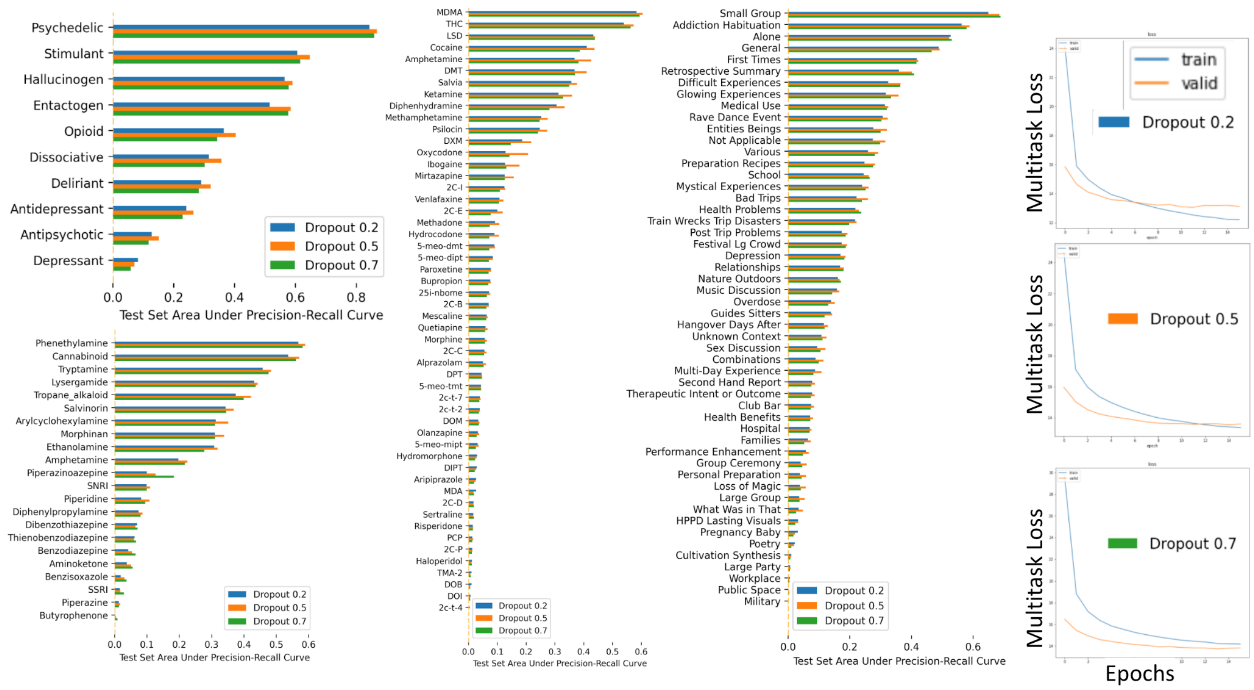

### Supplementary Figure 9: Tagscapes

This figure shows the relative ranking of tag predictions from BERTowid for a selection of key drugs. They are ordered separately for each drug, such that the tags at the top of the list, with blue bars are the most associated and tags lower in the list with red bars are progressively less associated with the given drug.

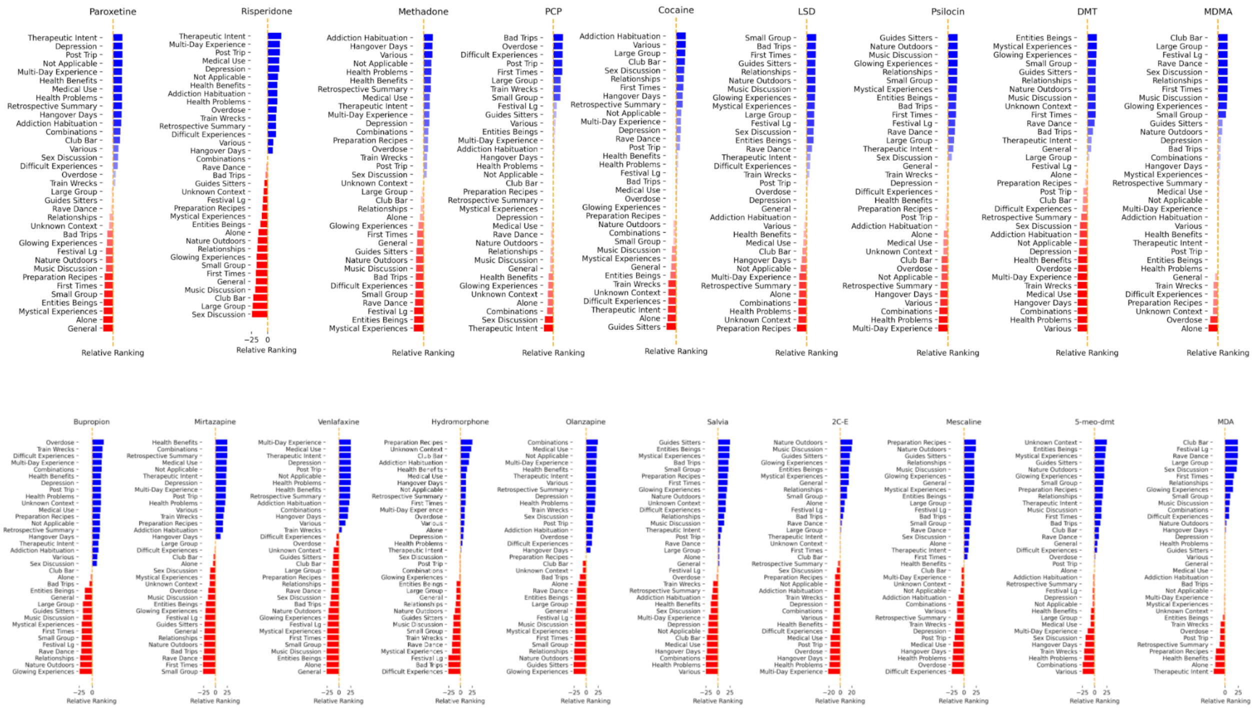

### Supplementary Figure 10: Receptor Affinities

Transformed receptor affinities for the 52 drugs and 61 receptors in the study, sourced from [37,39]. Affinity values are restricted to experiments with human proteins and aggregated in a hot-ligand sensitive manner. CCA was used to chart patterns of correlation between word-usage frequencies in drug testimonials and receptor affinity profiles.

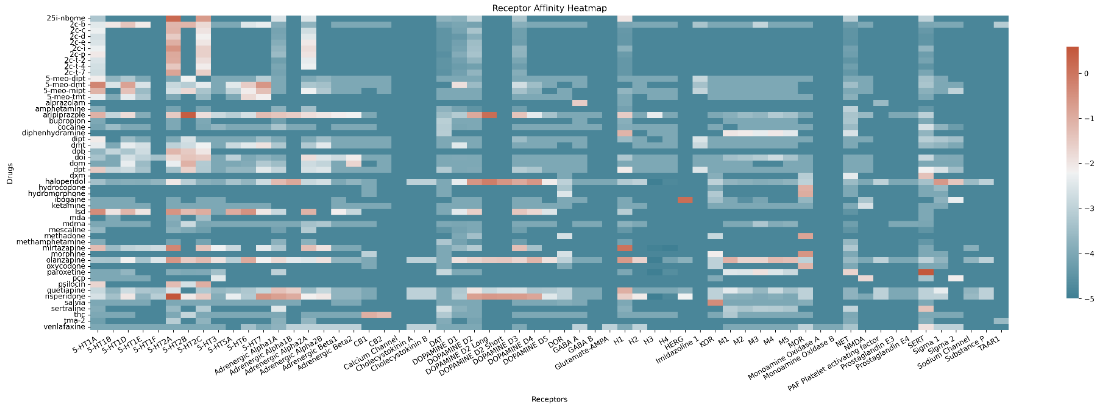

### Supplementary Figure 11: Tagscapes and Emotionscapes for DMT and MDMA

Panel 1: The first and last column show BERTowid’s tag predictions averaged for a given drug and ranked against all other drugs. The two middle plots show the analogous ranking from the 28 output heads of BERTiment.  BERTowid ranks DMT highly on “Entities/Beings” and “Mystical Experiences”, while BERTiment ranks it highly for “Surprise”, “Realization” and “Curiosity”. BERTiment’s MDMA rankings shows strong correlation with the hedonic-tone ordering and BERTowid ranks MDMA least likely to be tagged “Alone”.

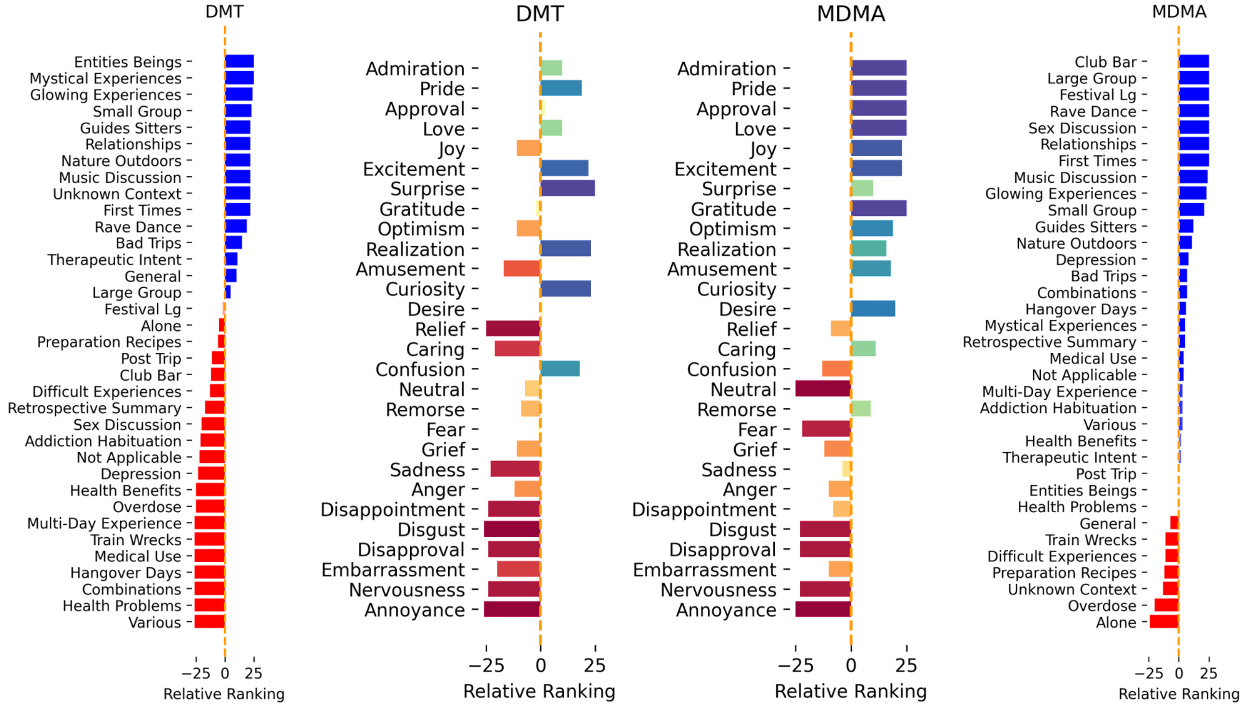

### Supplementary Figure 12: Tagscapes and Emotionscapes

Top, relative ranking of BERTiment predictions for drugs belonging to each of the 10 pharmacologic classes are shown, ordered by the hedonic tone spectrum derived from IMDB. Bottom, relative ranking of BERTowid tag predictions for the drugs belonging to each of the 10 pharmacologic classes.

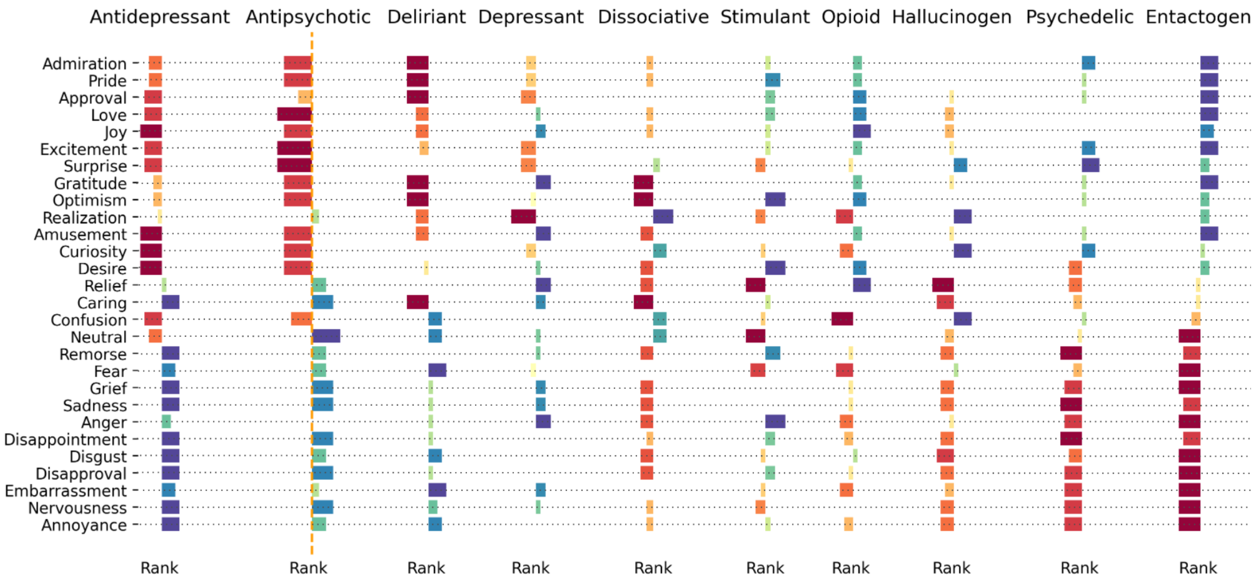


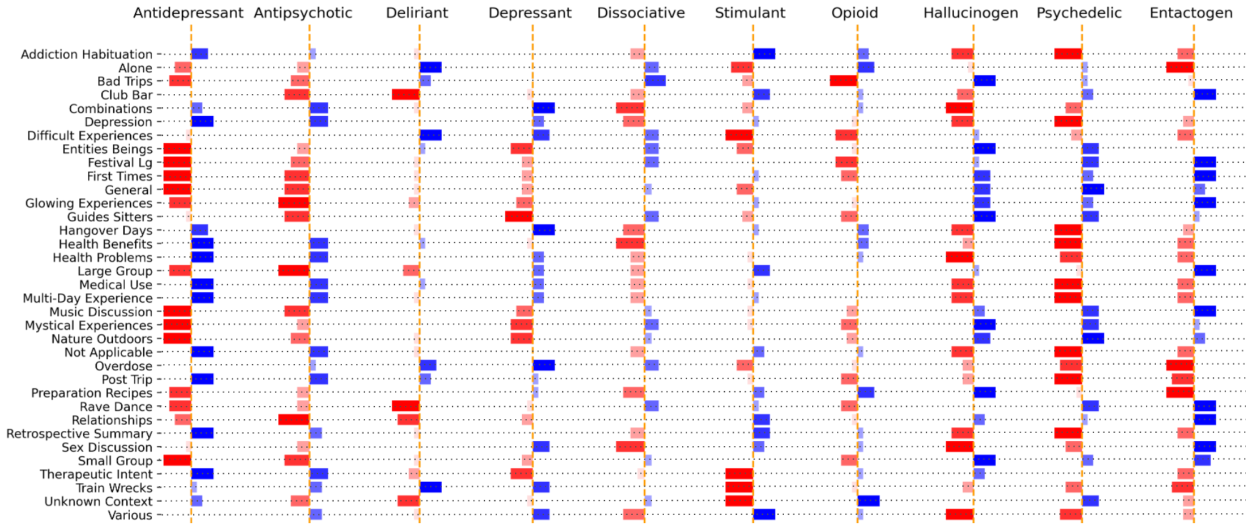

### Supplementary Figure 13: DTWs

Panel 1: Top left, BERTowid tag predictions for MDMA compared to all other drugs, ordered from most associated (blue bars) to least associated (red bars). Top middle, BERTiment sentiment averages compared all other drugs. Top right, DTW for “Love” showing MDMA’s prominence. Below, BERTowid tag DTW shows the universally highest ranking is for MDMA against all other drugs for tags “Club Bar”, “Rave Dance”, “Sex Discussions”, “Large Group”, and “Festival Crowd”.

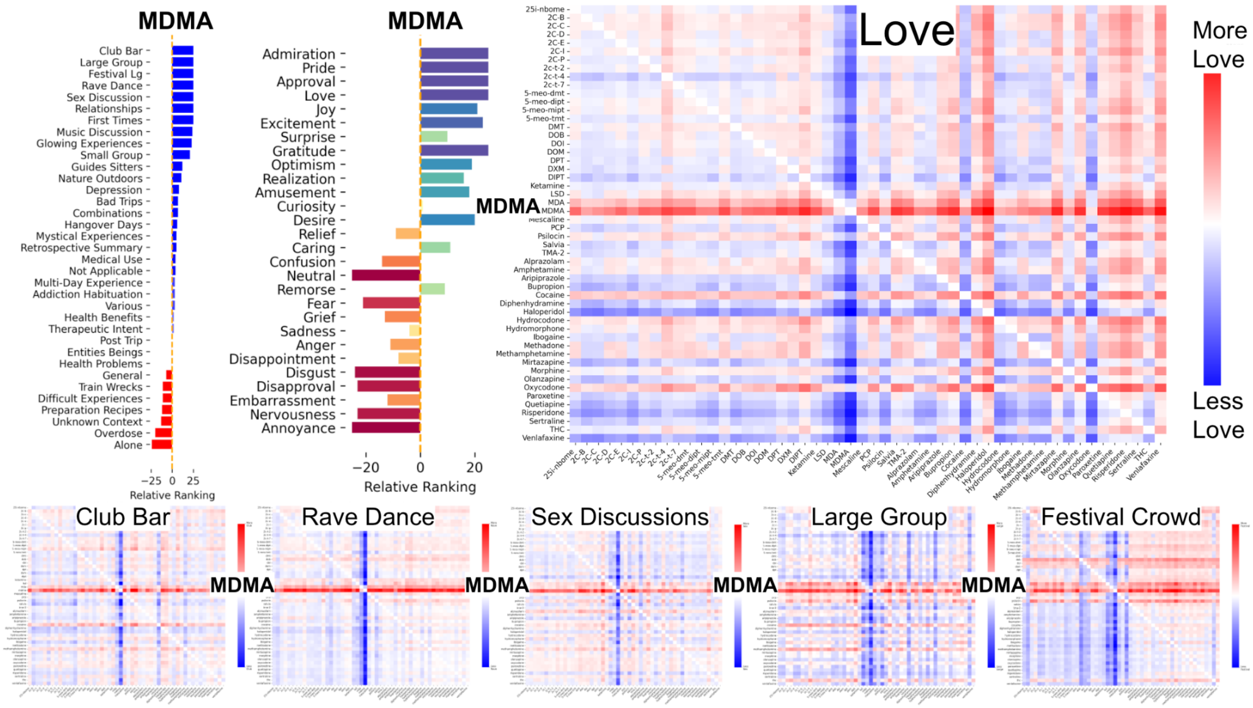

### Supplementary Figure 14: MDMA and Hedonic Tone

A subset of drugs representing the main pharmacologic classes are shown in terms of sentiment trajectories for “Love”, “Approval”, “Admiration”, “Pride”, “Joy”, “Excitement” (for which MDMA has high overall relative value throughout trajectory), as well as “Anger”, “Annoyance”, and “Disgust” (for which MDMA has low overall relative value through trajectory of drug experience). Lastly, the blue pole from CCA 5 reproduced below clearly highlights MDMA and related terms.

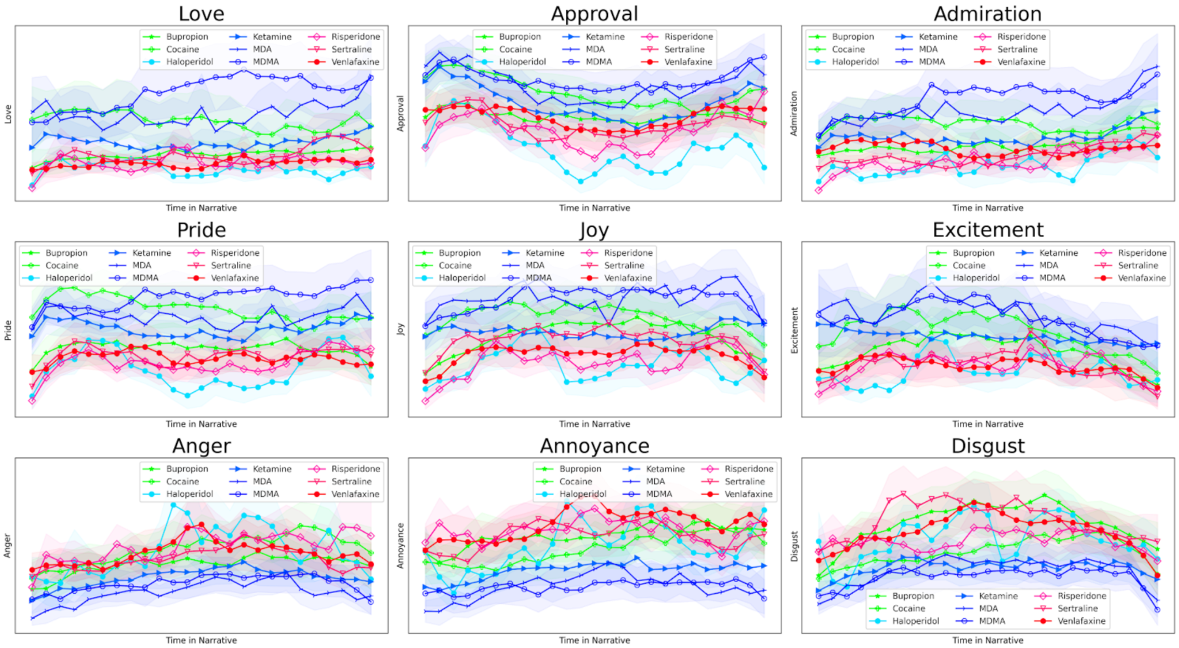


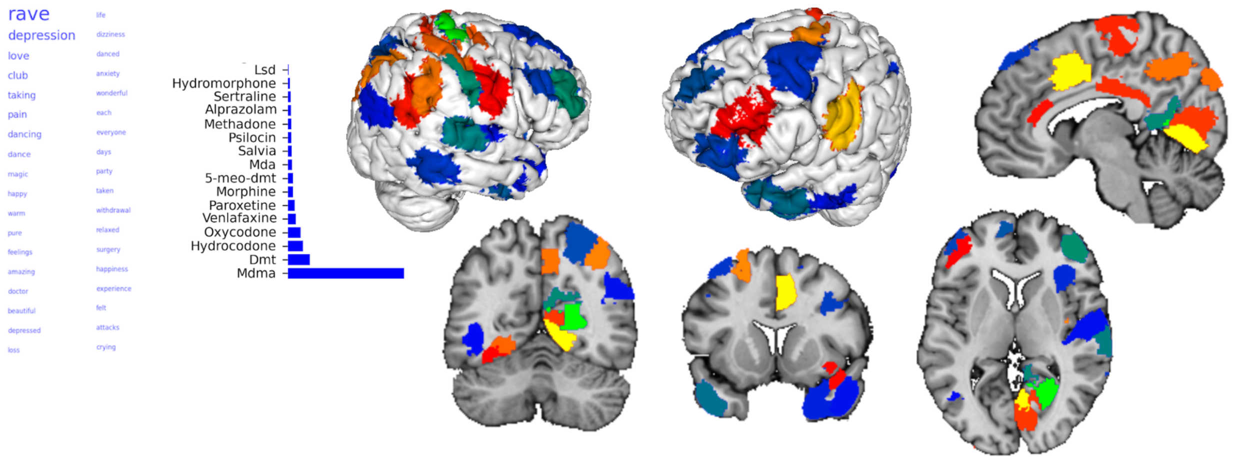

### Supplementary Figure 15: Trajectories of Mystical Experiences and Entities Beings

Comparison of all 52 drugs BERTowid tag trajectories for “Mystical Experiences” and “Entities Beings”.  Note the prominence of DMT and Salvia and how these tags, which are of great clinical interest, seem to highlight drugs from distinct pharmacological classes.

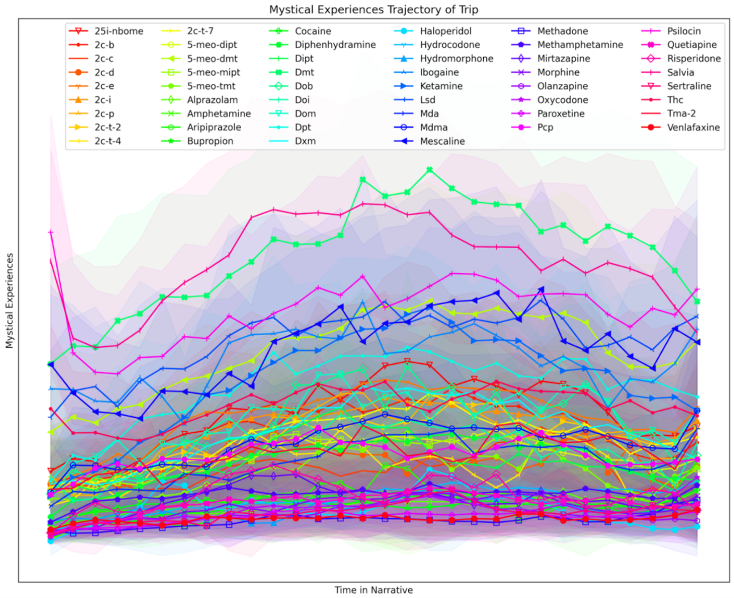


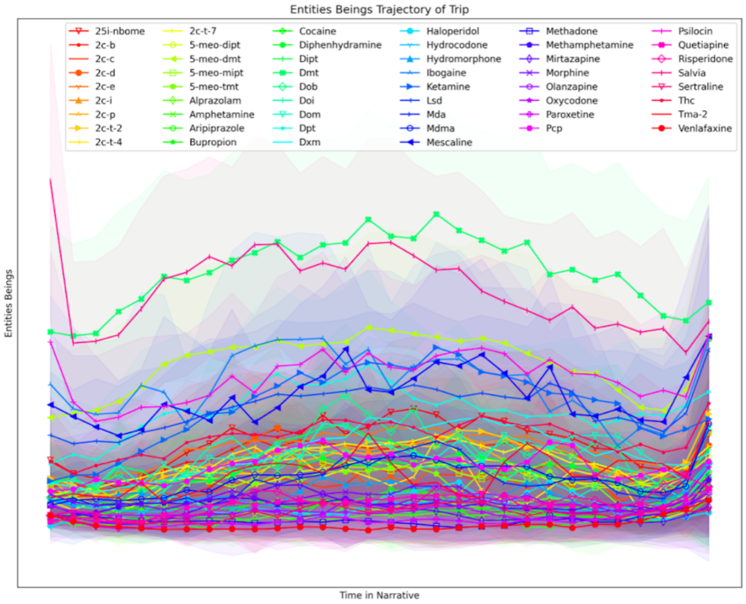

#### Supplementary Figure 16: Single Task vs Multi Task BERTowid

Comparing single-task or multitask BERT transformer models for BERTowid, it was found that they had comparable performance across drug pharmacologic classes as demonstrated in the AUC for Precision-Recall chart below; given the minimal difference, a multi-task BERT was used for drug class prediction. For sex prediction…

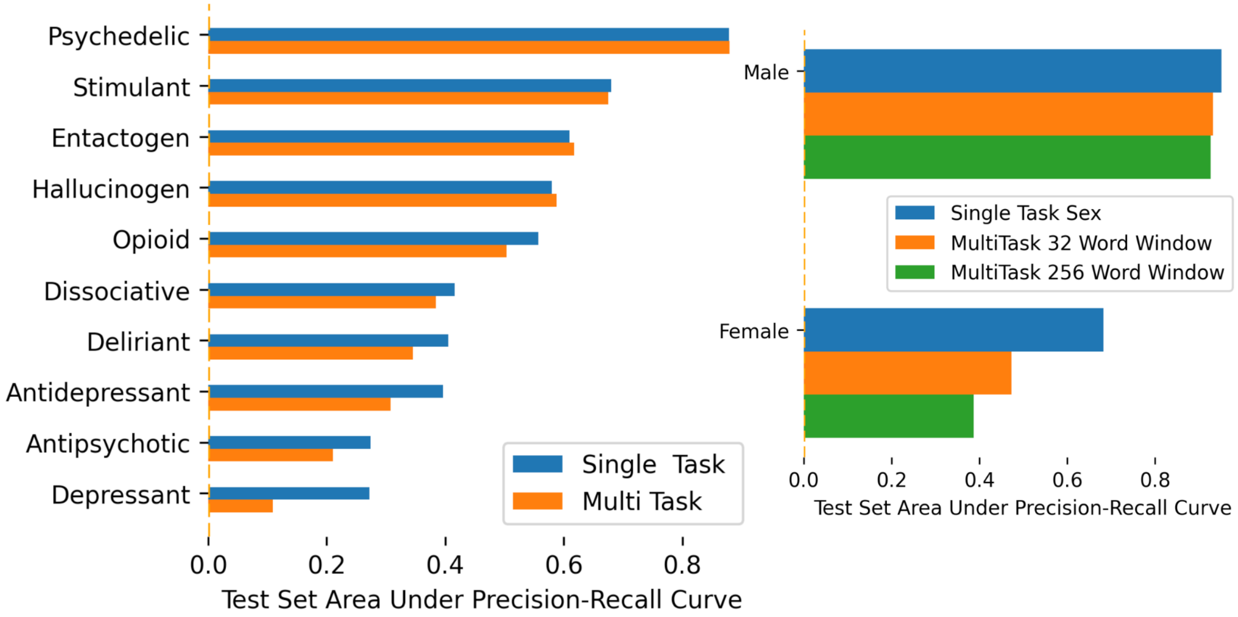

### Supplementary Figure 17: Cross Entropy Class Imbalance Weighting

Many of the categorical labels are class-imbalanced. This imbalance leads to poor performance on the less well-represented drugs and tags. To mitigate this we considered a weighted cross entropy loss, which scaled the loss by the inverse of each labels' prevalence to compensate for the imbalance. The weighted loss did increase precision for less well represented drugs (at the cost of reduced precision on the more prevalent drugs).  However, the results for tags were less convincing with only minor improvements in precision for the 4 rarest tags. A likely explanation is that less common tags are in fact less informative and may be applied less rigorously or consistently by the Erowid moderators.  Giving these less informative labels more weight in the loss function results in a worse model.

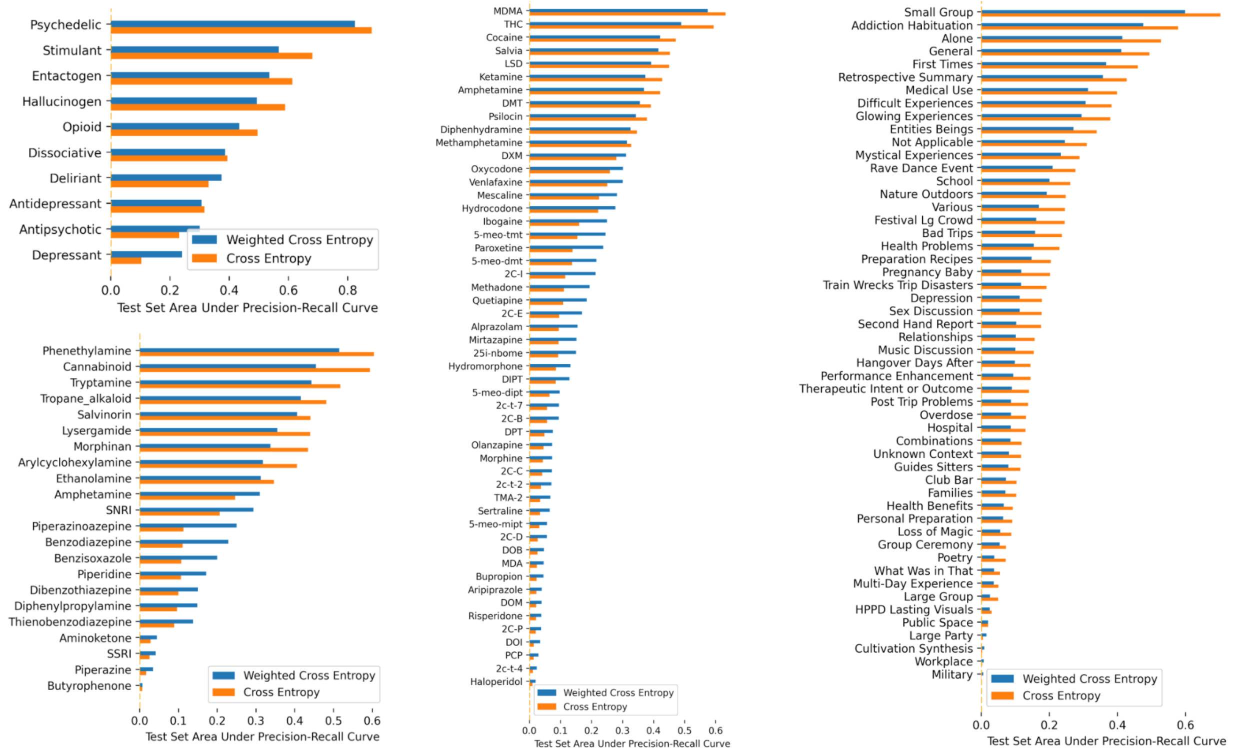

### Supplementary Figure: CCA 0

The dominant component across all drug experiences highlights a tension between terms relating to *lucid-beauty* and *mundane-suffering.* The most statistically significant CCA component linking receptor binding affinity to word usage across [x] words in11,816 testimonials of real-world experiences induced by 52 compounds is shown here. **Left**: A weighted list of words comprising the semantic theme of each extreme (red and blue) of this receptor-experience vector. Bigger words represent relatively larger ranking in this component. **Right**: Receptor affinity values most closely associated with this receptor-experience component are sorted by their relative ranking; blue bars correspond to blue words on the left while red bars correspond to red words. **Middle**: The middle panel shows how strongly each drug correlated to this receptor-experience component. Again, blue bars correspond to the blue words and receptors, while red bars correspond to the red words and receptors. **Main**: Brain renderings and surfaces highlight which of the 200 parcellated anatomical regions were found to most densely express the receptor-gene RNA as described by Allen's Human Brain Atlas. Sagittal left, sagittal right, and axial brain slices are shown at x = -7, y = +8, and z = +8 of MNI-space, respectively. The colors of each rendering represent the density of weighted receptor gene expression for each region and correspond to the extremes of this component: blue words and bars correspond to blue/greed brain regions (green > blue intensity), while red words and bars correspond to the red/yellow brain regions (yellow > red intensity). The surfaces show the weighted receptor expression on the surface of the cortex using Mango.

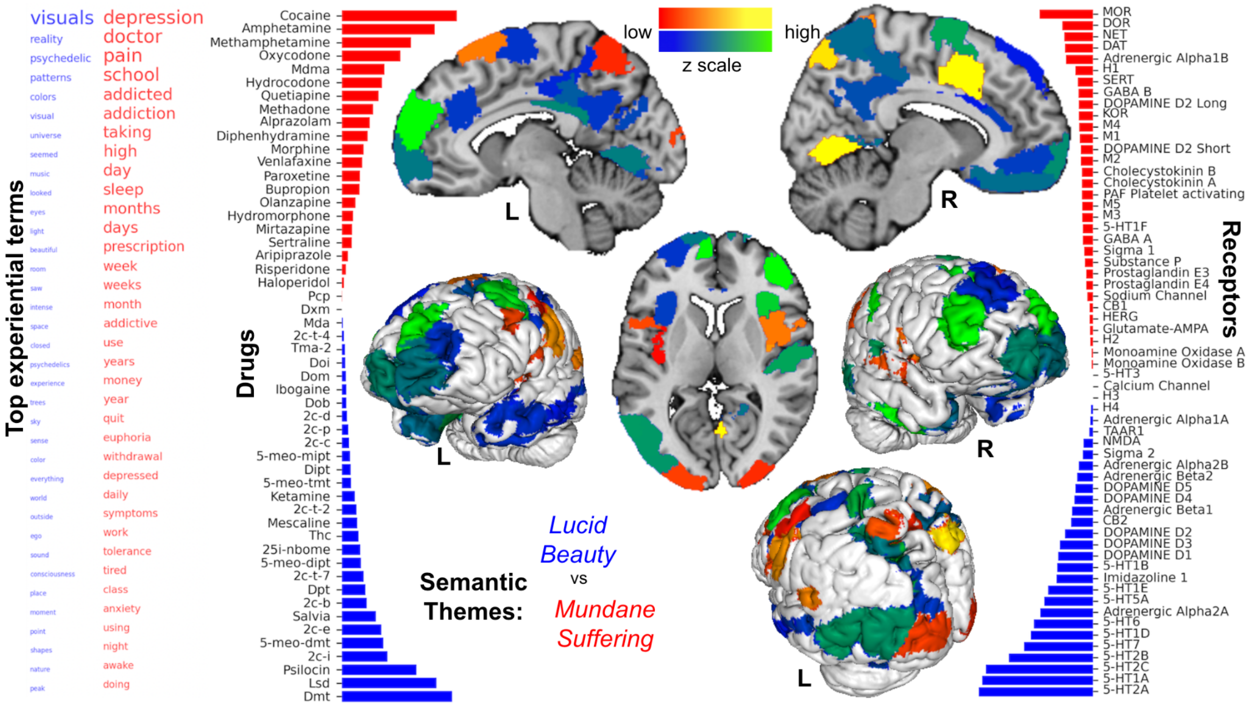

#### Supplementary Figure: CCA 1

The second significant component underlying drug experience charts a dichotomy between *depression-insomnia* and *impulsivity-addiction.* This component captured a motif of impulsivity (*rush, party, craving)* and addiction (*dealer, addicted, hooked, bought*), see Supplementary Figure: CCA 1. These terms were associated with the drugs cocaine, methamphetamine, and MDMA, as well as receptor affinity at CB1, DAT, CB2 and with expression of these receptor genes in the primary visual and sensory cortices. The opposite extreme flagged terms relating broadly to mental illness (*anxiety, depression, symptoms, diagnosed, psychotic, bipolar*) and treatment (*doctor, psychiatrist, medicine, treatment)*. This semantic theme was associated most with the drugs quetiapine, diphenhydramine, and DXM, as well as receptor affinity to H1, 5-HT2A, and D2 receptors. The expression of these receptor genes was found to be most dense across the distribution of the default mode network including the posterior cingulate cortex, ventromedial prefrontal cortex, and the inferior parietal lobules. Sagittal and axial brain slices are shown at x = 5 and z = -11.

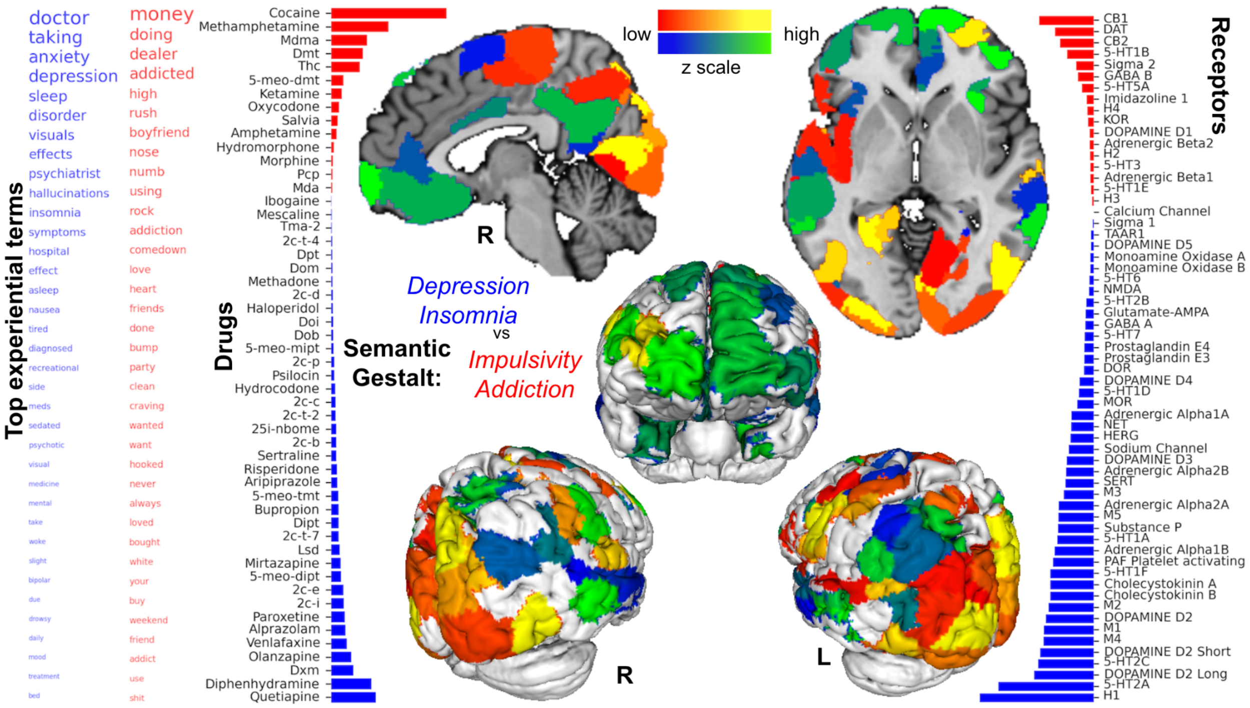

### Supplementary Figure: CCA 2

The third significant receptor-experience component tracked two extremes: *perception-celebration* and *cosmic-expansion*. This component identified, at one extreme, a collection of terms denoting cosmic (*universe, space, dimension, breakthrough*), awareness (*remember, dream, consciousness*) qualities of subjective experience, and entities (*beings, spirit, entity, alien*) that were nested in a temporal horizon of immediacy (*seconds*). This conceptual cluster was associated with drugs THC, DMT, Salvia, Ketamine, and 5-MeO-DMT, as well as receptor affinity at CB1, CB2, KOR, M1, and 5-HT7, and the expression of these receptor genes was most dense in the medial prefrontal cortex, dorsal anterior cingulate cortex, and temporoparietal junction. The opposite extreme of this component was characterized by terms relating to perception (*visuals, music, tracers*), somatic sensation (*nausea, effects, stomach, ingested, sex*), and intensity (*peak, comedown, amazing, mild*) situated in a time-horizon of *hours.* This aggregation of terms was associated with drugs MDMA, 2C-I, and 2C-E, as well as receptor affinities at 5-HT2A, Alpha 2A, and 5-HT2C, and the expression of these receptor genes was most dense in the primary visual cortex as well as areas of the primary sensory and motor cortices. Sagittal and axial brain renderings are shown at x = -6 and z = 8.

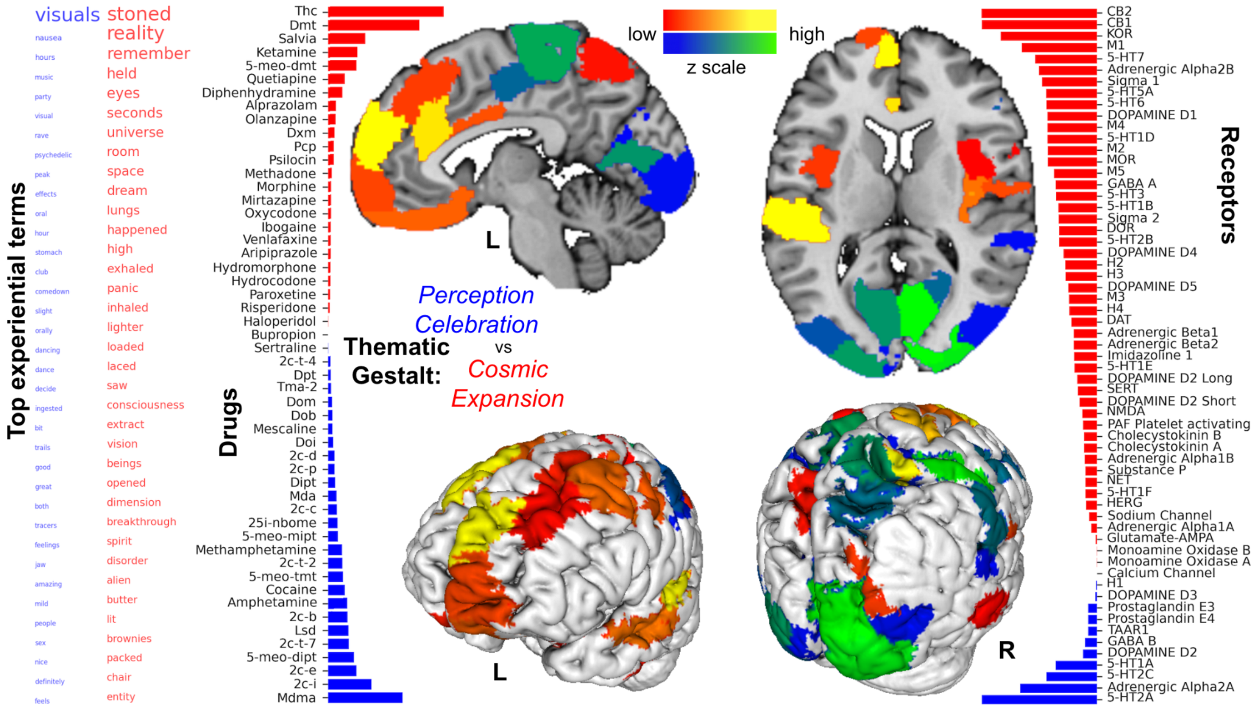

### Supplementary Figure: CCA 3

Receptor-experience component CCA 3 highlighted *bad-high* and *pain-relief.* This component tied together a range of terms relating to psychopathology including (*depression, anxiety, panic, paranoid, insomnia, psychotic*), diagnosis (*psychiatrist, disorder, diagnosed, symptoms*), and substance use (*stoned, rave, party, club, laced*). This theme of substance-related psychopathology was associated with drugs THC, MDMA, LSD, methamphetamine, and amphetamine. Receptor affinity weightings favored 5-HT2B, Alpha 2A, and Alpha 1A, while gene receptor expression was most dense in the medial prefrontal cortex, anterior insula, and dorsal anterior cingulate cortex. The opposite pole of this vector charted a subjective theme of discomfort (*pain, nausea, itching, sick)* and comfort (*warm, relaxed, euphoria, nice)*. This semantic spectrum of comfort was associated with drugs oxycodone, hydrocodone, DMT, methadone, and ketamine, receptor affinity for MOR, KOR, and DOR receptors, and receptor gene expression in the primary visual cortex, precuneus, and mid-cingulate cortex.Sagittal left, sagittal right, and axial brain renderings are shown at x = -6, x = +8, and z = -9, respectively.

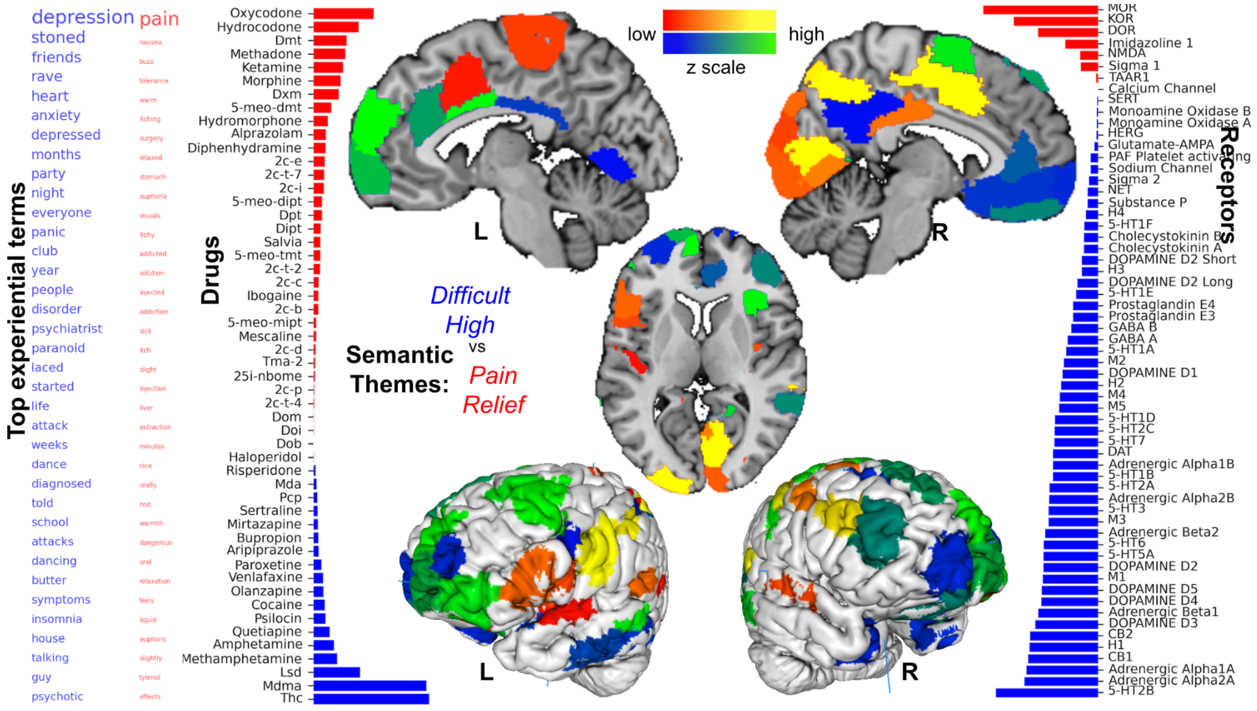

### Supplementary Figure: CCA 4

The fifth receptor-experience component uncovered *edibles* and *bipolar* phenomena. This component identified by CCA captured a semantic theme rich with terms relating to both the manic *(energy, insomnia, rush, intensity, psychotic, money*) and the depressive *(depression, mood, insomnia, anxiety)* phases of bipolar illness. This theme was associated with drugs DMT, cocaine, 5-MeO-DMT, and methamphetamine, as well as receptor affinity at 5HT1A, 5HT7, 5HT2C, and 5HT1D, and the expression of these receptor genes was densest across the distribution of the default mode work (medial prefrontal cortex, posterior cingulate cortex, medial temporal lobes) and salience network (dorsal anterior cingulate cortex, anterior insula). The converse semantic extreme of this component clustered words relating to inebriation (*stoned, high, laughing*), somatic discomfort (*cough, pain, itching, sick*), and oral consumption (*laced, brownies, food, butter, bowls, bottles, drank*). This linguistic gestalt was linked to drugs THC, DXM, and diphenhydramine, receptor affinity at CB1, CB2, and MOR, and receptor gene expression especially in the visual cortex. Sagittal left, sagittal right brain renderings are shown at x = -6 and x = +8, respectively.

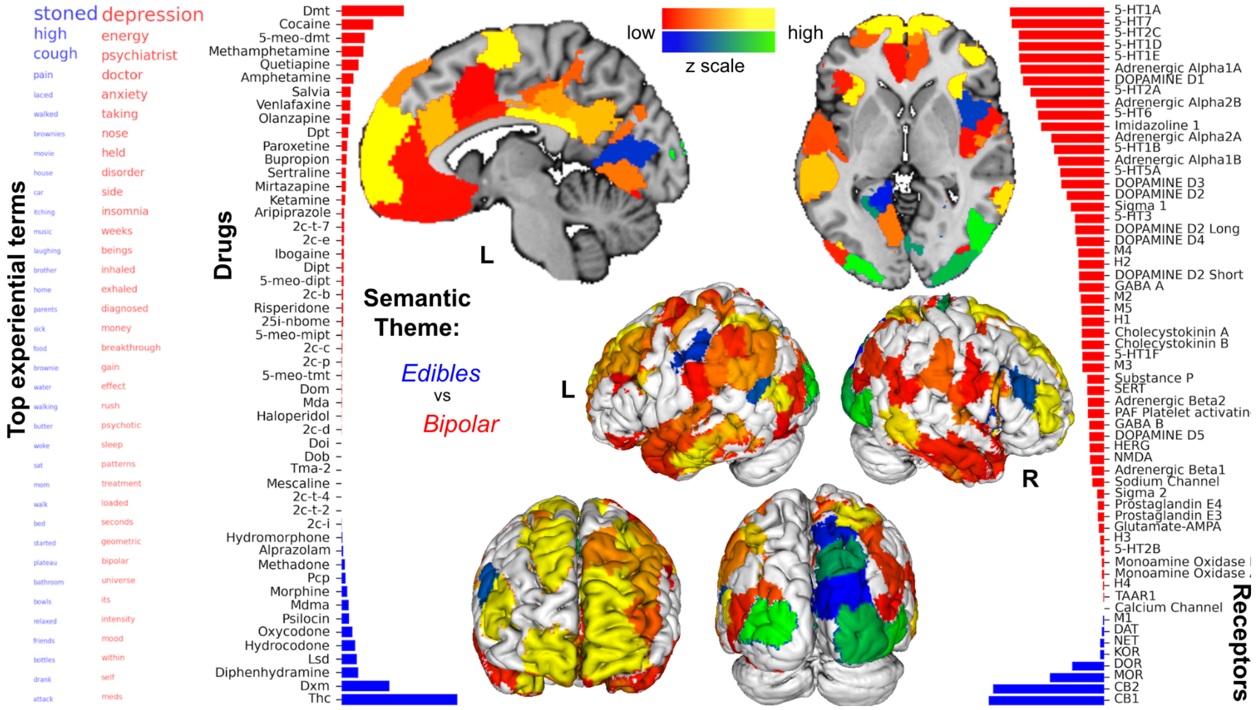

### Supplementary Figure: CCA 5

The sixth receptor experience component revealed *emotional-extremes* and *drug-induced-psychosis.* This component identified a constellation of terms describing a rich array of emotions, ranging from positive (*love, happy, warm, pure, amazing, beautiful, wonderful, relaxed*) to negative (*depression, loss, anxiety, crying*), as well as a context of a nightclub or “rave” (*rave, club, dancing, party*). This semantic pattern was strongly linked to drug MDMA, with receptor affinity at 5HT2B and Alpha 2B, and receptor gene expression isolated to the primary visual cortex and primary sensory regions innervating the trunk, head, and neck. The opposite extreme of this component flagged a pattern of terms tying together psychosis (*visuals, hallucinations, auditory, voices, hospital, psychotic, psychosis*) and addiction (*money, bump, dealer, addiction, addicted, addict*). This semantic theme was associated with receptor affinity at H1, 5-HT2A, and CB2, as well as receptor gene expression in the mid-cingulate and visual cortices. Sagittal left, axial, coronal posterior, and coronal anterior rendered brain slices are shown at x = +8, z = +4, y = -57, and y = +6, respectively.

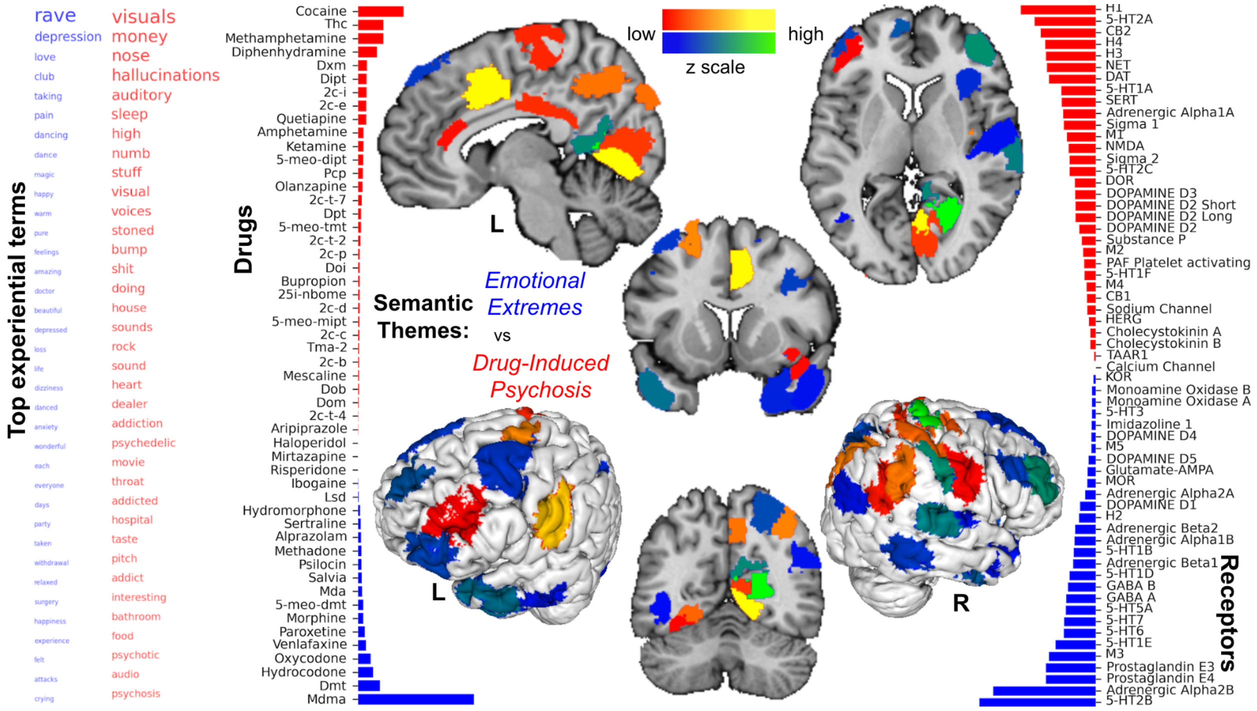

### Supplementary Figure: CCA 6

Receptor experience component CCA 6 revealed *emotional-extremes* and *drug-induced-psychosis.* This component distilled a semantic cluster of terms relating to somatic sensation (*pain, nausea, ate, warm, heart, relaxed*) and intoxication (*stoned, high, laced, strong, brownies*). This constellation of terms was mostly linked to THC, as well as receptor affinity at 5-HT2A, CB2, and CB1, and receptor gene expression isolated to the primary sensory cortex corresponding to the dermatomes of the head, neck, and torso. The opposite extreme of this component flagged words relating to liminal states (*plateau, remember, drunk, hallucinations, woke, spiders, night, memory, hole*). This semantic theme was associated with drugs DXM, MDMA, diphenhydramine, and ketamine, as well as receptor affinity at SERT, NET, and M3 receptors with gene expression for these receptor subtypes isolated primarily to the mid-cingulate cortex.  Sagittal left, sagittal right brain renderings are shown at x = -6 and x = +8, respectively.

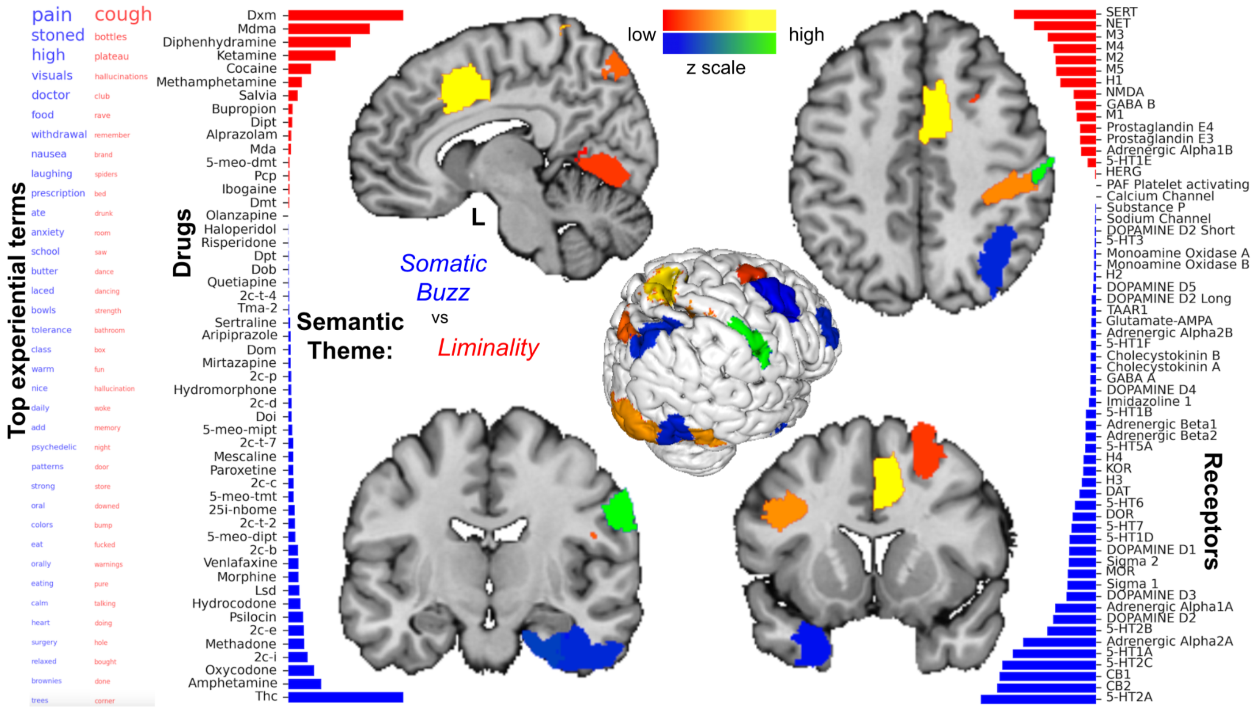

### Supplementary Figure: CCA 7

Receptor-experience component CCA 7 highlighted *attention* and *auditory-distress.* This component also identified terms describing audition (*auditory, sound, pitch, voices, audio*) and distress (*depression, withdrawal, anxiety, suffered, suicidal, panic*). These terms were associated with drugs amphetamine, DiPT, venlafaxine, paroxetine, as well as receptor affinity at SERT, NET, CB2, Sigma 2, and expression of these receptor genes in the salience network (dorsal anterior cingulate cortex and anterior insula) and medial prefrontal cortex. The opposing extreme linked a set of terms relating to psychosis (*psychotic, psychosis, visuals, hospital*) as well as various levels of arousal and hedonic tone (*sedated, lethargic, tired, pleasant, amazing, beautiful, wonderful*). This semantic pattern was associated with drugs quetiapine, MDMA, LSD, and olanzapine, as well as receptor affinity at 5-HT2A, 5-HT2C, D3, and D2 receptors, and receptor gene expression in the visual and mid-cingulate cortices.  Sagittal, axial ventral, axial dorsal, and coronal brain slice renderings are shown at x = -5x, z = -8, z = +18, and y = -6, respectively.

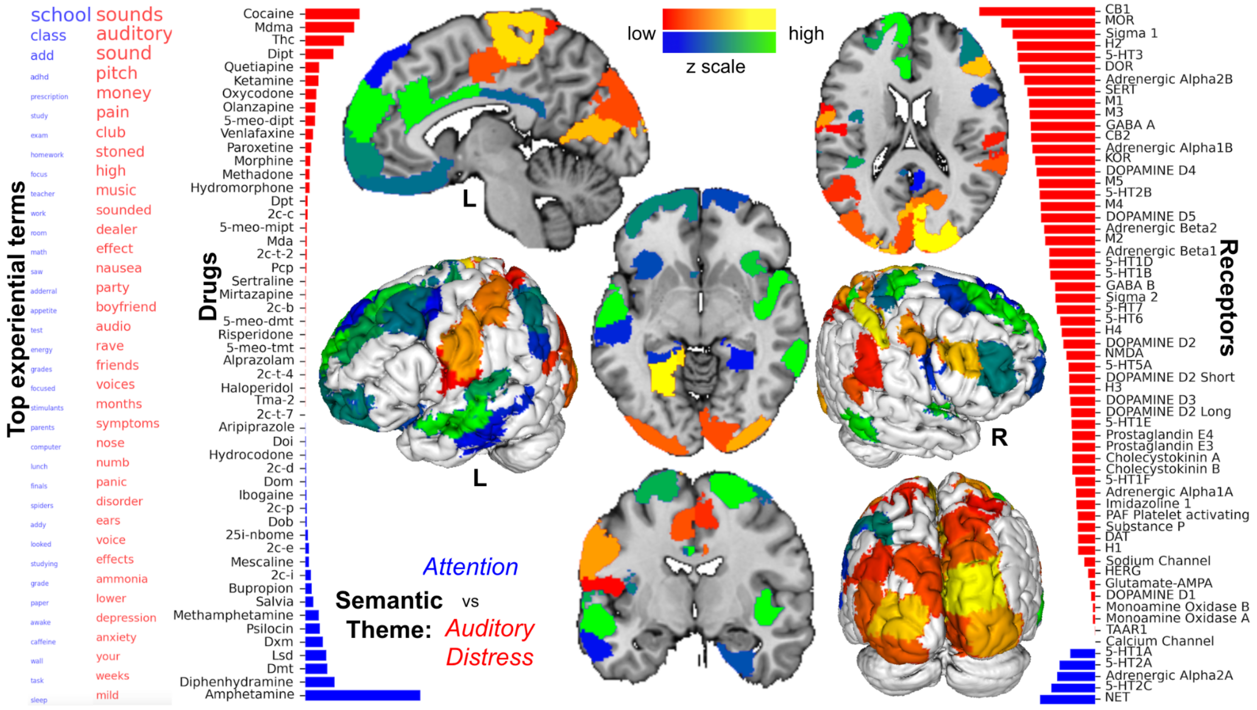

### Supplementary Figure: CCA 8

This component identified a similar set of terms describing audition (*auditory, sound, pitch, voices, audio*) and distress (*depression, withdrawal, anxiety, suffered, suicidal, panic*). These terms were associated with drugs amphetamine, DiPT, venlafaxine, paroxetine, as well as receptor affinity at SERT, NET, CB2, Sigma 2, and expression of these receptor genes in the salience network (dorsal anterior cingulate cortex and anterior insula) and medial prefrontal cortex. The opposing extreme linked a set of terms relating to psychosis (*psychotic, psychosis, visuals, hospital*) as well as various levels of arousal and hedonic tone (*sedated, lethargic, tired, pleasant, amazing, beautiful, wonderful*). This semantic pattern was associated with drugs quetiapine, MDMA, LSD, and olanzapine, as well as receptor affinity at 5-HT2A, 5-HT2C, D3, and D2 receptors, and receptor gene expression in the visual and mid-cingulate cortices. Sagittal, axial ventral, axial dorsal, and coronal brain slice renderings are shown at x = -5x, z = -8, z = +18, and y = -6, respectively.

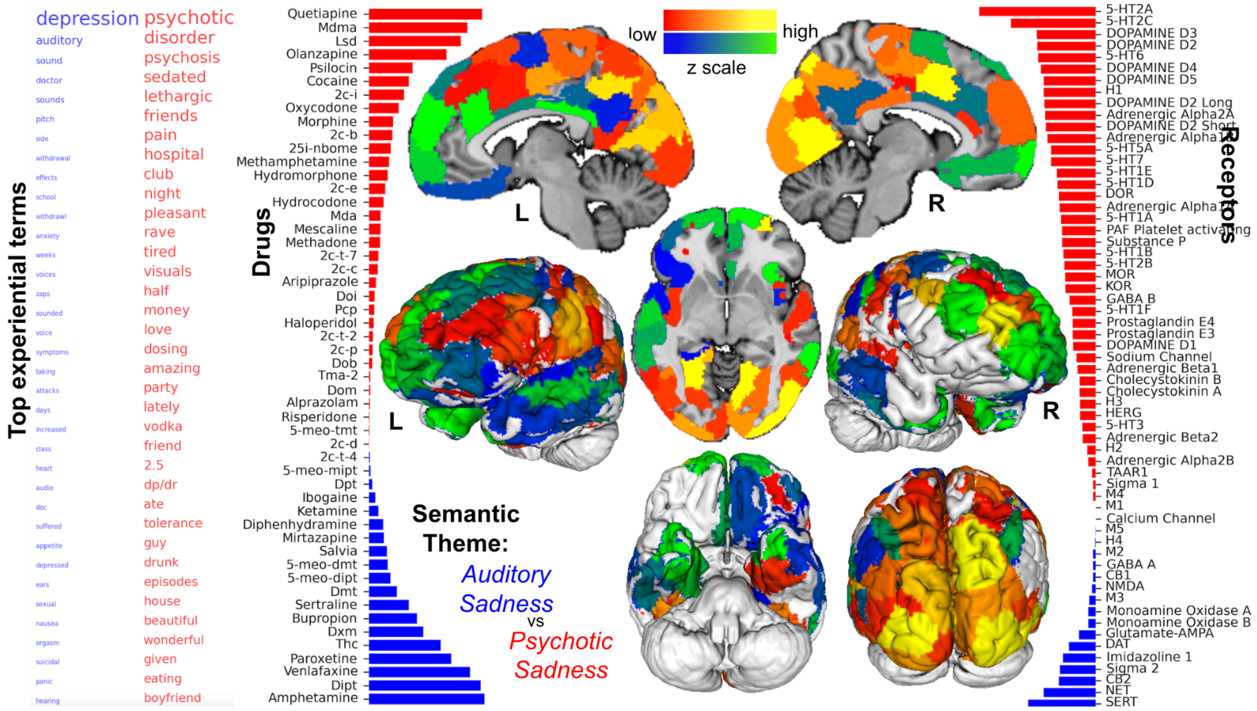

### Supplementary Figure: CCA 9

Receptor-experience component CCA 9, shows *sexual-angst* and *auditory-inensity.* This component highlighted a semantic theme angst (*depression, anxiety, attacks, panic, suicidal, depressive, sick*) and sexuality (*sexual, orgasm, sex*). This semantic pattern was associated with drugs DXM, cocaine, venlafaxine, and paroxetine, as well as receptor affinity at SERT, 5-HT2A, and 5-HT2C, and gene expression of these receptors in the visual and primary sensory cortices. The opposite extreme of this component tracked a set of terms relating to sound (*auditory, sound, voices, pitch, audio, music, ringing*) and arousal (*rave, love, buzz, crash, dance*). This semantic gestalt was associated with drugs amphetamine, MDMA, DiPT, and DMT, as well as receptor affinity at 5-HT2B, Alpha 2B, imidazoline 1, and CB1. Expression of these receptor genes was most dense in the primary and secondary auditory cortices, anterior insula, dorsal anterior cingulate cortex, and medial prefrontal cortex. Sagittal left, sagittal right, axial, and coronal brain slice renderings are shown at x = -6, x = +41, z = -3, and y = +25, respectively.

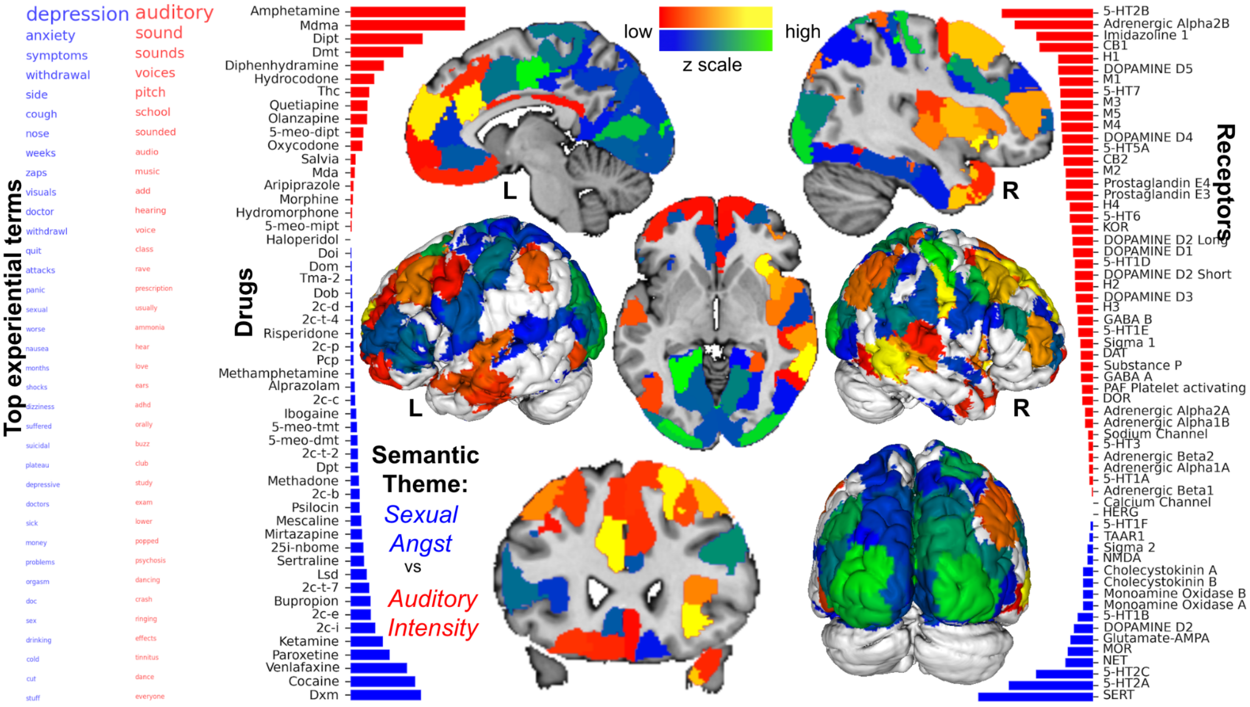

### Supplementary Figure: CCA 10

Receptor-experience component CCA 10 shows *sexual-angst* and *auditory-inensity.*
*This component tracked a semantic constellation of terms relating to retching (cough, pukes, nausea, stomach, vomit) and ailment (itching, sick, doctor, diagnosed, psychiatrist, pain, warnings, meds, disorder). This phenomenal pattern was most associated with drug DXM and amphetamine, as well as receptor affinity at SERT, MOR, KOR, and DOR, and with expression of these receptor genes in the primary visual cortex, mid-cingulate cortex, and bilateral posterior insula. The opposite extreme of this component identified a set of words evoking eerieness (spiders, hallucinations, weird, freaked, anxiety) and physical environment (mom, car, door, wall, room, home, driving, corner, fell). This final semantic pattern was associated with drug diphenhydramine, LSD, and alprazolam, as well as receptor affinity at H1, 5-HT2A, and 5-HT2C, and with expression of receptor genes in the precuneus and primary sensory cortices.*
Sagittal, and coronal brain slice renderings are shown at x = +7, z = 0, and y = +1, respectively.

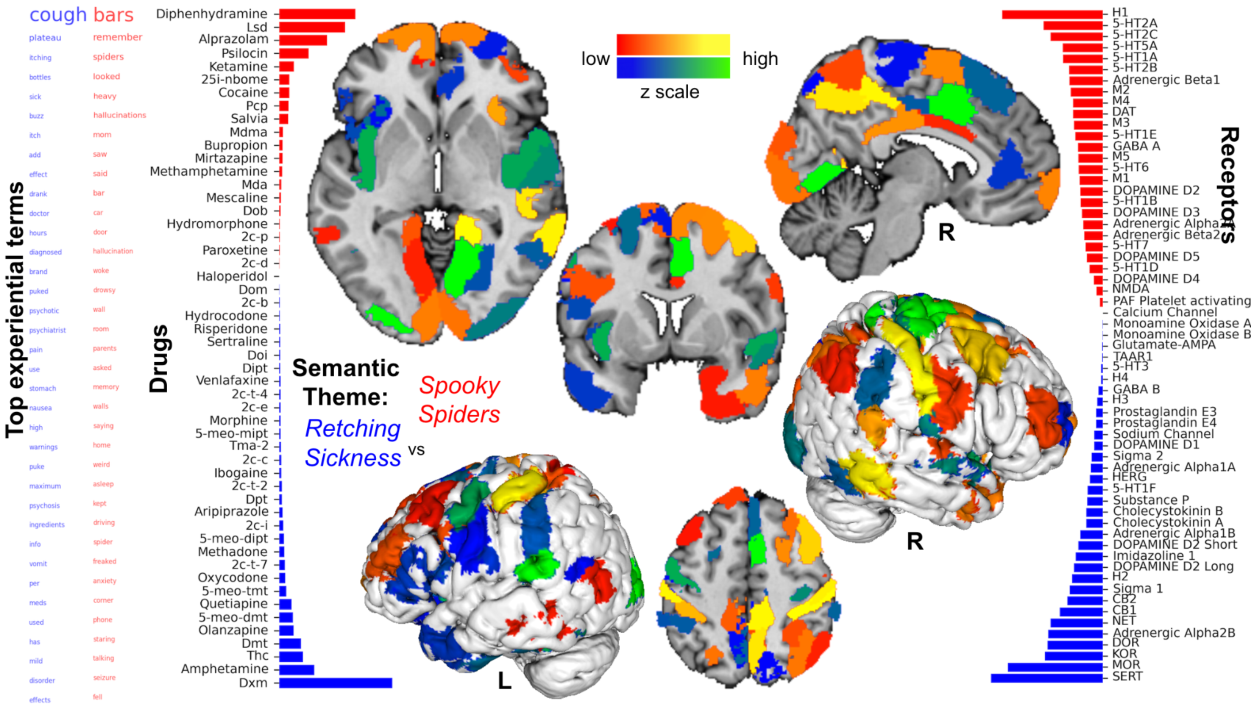

#### Supplementary Table 1: Extreme BERTiment

Below are the highest and lowest ranking ~32 word testimonial-windows for each of the 28 sentiments. The unmistakable clarity of sentiment contained in both the sentiment label excerpts and also their opposites suggest that BERTiment is describing meaningful sentimental valences in both the positive and negative directions for all 28 dimensions of analysis.

| sentiment | drug | text |
|---|---|---|
| admiration | 2c-e | It was amazing and incredibly powerful. And the visuals continued to grow in scope and magnitude seemingly every minute. Eventually, everything became in constant motion and all objects were continuously undulating and before our very eyes. |
| not_admiration | 2c-t-4 | I didn't know where I was or how I'd gotten there. I didn't remember that I'd taken DRUG and I didn't remember anything that'd happened earlier. Somehow I got it in my head that I'd died and was in a waiting room, waiting for my soul be judged and sent to either Heaven or Hell. |
| amusement | 25i-nbome | At one point I felt it was as if Jesus telling me I turned on him. But then I laughed at him and the idea. No offense to any christians reading this. I don't hate jesus, it was just a crazy trip lol. |
| not_amusemen t | haloperidol | There are other effects I've been noticing lately, perception-wise, but it is extremely difficult to describe them with words. These include psychological changes, namely a more objective perception of feelings and emotions in one self and other people, which is both positive and negative since it involves a degree of detachment from people, which some might consider dangerous or conflictive. |
| anger | thc | How DARE he. What the FUCK was he thinking – not telling me what the fuck I was smoking. I was going to tell him off the next time I saw him – no – punch the bastard out – kill him? |
| not_anger | risperidone | Of all the many substances I've consumed, Matrix is still my favorite. |
| annoyance | amphetami ne | When I'm disturbed while I'm trying to get work done on DRUG I immediately want to tell that person off so they'll leave me alone and let me work. If my concentration is broken, it is really annoying. |
| not_annoyance | risperidone | The following day, I was invited by my friend to a concert in our area, at a large amphitheater, with a well-known band as the headliner. I was a musician in a band for many years and I've attended quite a few shows, including a couple at that same venue, so I was excited and looking forward to this day. |
| approval | 5-meo-dmt | I agree, but I would start with tasters when I was sober if I'd never been there before. the Teafærie |
| not_approval | tma-2 | What day of the week was it? How does a day fit into a week? A week into a month? A month into a year? A year into a century? A century into history? |
| caring | amphetami ne | Stay safe friends! |
| not_caring | aripiprazol e | Maybe it's the reptilian brain exercising it's serpentine nature... I'm not sure at this point. I turn on my TV at 9:00 p.m. EST, Cartoon Network, Scooby Doo, one of the movies too. Can't remember which one, but I find cartoons in this state VERY startling. |
| confusion | thc | Eventually I started becoming very confused. All of the music they were playing in the background was really standing out to me and I had no idea what the episode was about. Hell, I couldn't even tell what the jokes were. |
| not_confusion | tma-2 | Happy travels friends. |
| curiosity | dxm | DRUG DRUG DRUG Something to do with a Chandelier? I have had three or four previous DRUG experiences, but I wanted to try a higher dose so to get to the third plateau out of curiosity. |
| not_curiosity | haloperidol | Music is my canvas. |

| | | |
|---|---|---|
| desire | 2c-i | I just could not resist hunting some porn and pleasuring myself. I wish I could have shared the experience with a female. The sexual energy was so intense, I could vividly fantasize about what I wanted. |
| not_desire | risperidone | Increased muscular tension, especially prevalent in facial area, including tightness of the jaw. OEVs becoming slightly apparent. 11:00 (T+02:00) Heavy OEVs, full visual distortions and multitude of tracers. Sparkling of colors throughout visual field, especially reds greens and blues. |
| disappointment | 2c-t-7 | Still, the DRUG seemed to put me in a good mood and it wasn’t at all unpleasant to be in a club environment. The DJ stopped spinning at 1:30 a.m and I was disappointed that things were ending so early. |
| not_disappointment | risperidone | So I’ll leave it at that. This has been my shamanic initiation. It was a pleasure sharing it with you. As Terence McKenna said, a Shaman is someone who has seen the beginning and seen the end. |
| disapproval | bupropion | I wouldn't even suggest trying this drug, it doesn't seem to have any positive effects. |
| not_disapproval | venlafaxine | See you all there x |
| disgust | 5-meo-mipt | 20:30 – Eat ~7mg of DRUG (scale was fluctuating slightly between 6 and 7 milligrams). Tastes disgusting. For 7mg of DRUG this shit sure is strong. Tasted similar to DRUG but it was even more bitter. |
| not_disgust | haloperidol | In the morning, everyone who had participated in the ceremony visited with each other and shared some of their experience. I sought out Sharon to thank her for standing in for me and accepting Prem Das's gift. |
| embarrassment | psilocin | Embarrassed to be acting so nuts in front of my friends and not wanting to affect their trips, I turned to face the back wall and leaned forward onto the bed standing now while tightly grasping the sheets as the wave overtook me. |
| not_embarrassment | risperidone | Also highly interested in virola now! Deep thanks to those who made this long-wished-for experience manifest. |
| excitement | 2c-c | She told me to call her around 2:30 to let her know how I was doing. I can't wait for 2:30. I was so excited. The drug had already started to manifest itself. I felt the normal stomach clenching I do on every DRUG I have tried. |
| not_excitement | 2c-t-4 | This DRUG is grossly misunderstood and understudied. |
| fear | mda | go away. This frightened me extremely, as I was afraid of getting rabies if it attacked me. But I seemed detached from my fear and instead watched it for several minutes (if it was even there). |
| not_fear | haloperidol | Either way, I greet you with: Namaste. |
| gratitude | psilocin | “Thanks, for bringing me back to awareness of the others in this house. Thank you all, for existing, and not-existing with me, and having this experience. I mean, it’s really quite profound, don’t you think? |
| not_gratitude | haloperidol | For example, the ceiling would come apart in abnormally-shaped sections and each section would raise up or lowest slightly so that they were all at different heights, and then they would shift around to gently overlap each other. |
| grief | ketamine | I saw paramedic's put me in an ambulance and take me to the hospital, I saw my family and friends standing around me, and I saw my grave. I thought to myself 'So this is death. |
| not_grief | haloperidol | “Intrigued and Interested” is a better descriptor: moreso than other substances in the past. We each had two coffees (t-4:00), 8 keto mini pancakes for breakfast (t-2:00), one Rockstar each (t-1:00), plus our normal regimen of supplements. |
| joy | alprazolam | Euphoric and feeling great. Even nodding a little bit, but not the kind of nod that makes me want to go to sleep, just nod and enjoy it. 7:45 Really talkative and happy, feeling the effects of everything, mostly the DRUG though. |
| not_joy | mirtazapine | but it's not the material itself, what's dangerous, but the people who and how use it. |
| love | psilocin | I love you I love you I love you I murmur, saying it as the world to myself as myself to the world, begging for forgiveness and comforting with infinite love, a lover in passion a throwaway comment to a friend a parting kiss a greeting touch, I love you I love you. |
| not_love | doi | After several minutes of debating back and forth how easily I could hide my obviously impaired state, the man left; he turned out to simply be a cable television representative making a cold call. |
| nervousness | 5-meo-mipt | I was feeling very tight chested and uncomfortable, so I smoked a bowl of DRUG to take the edge off. I found my mind wandering quite a bit, and worrying about how the rest of the trip was going to progress considering how high I had already become. |

| | | |
|---|---|---|
| not_nervousness | haloperidol | He now has more of a respect for Salvinorin A. |
| optimism | methamphetamine | I didn't graduate high school, so I live off on disability. This hopefully concludes my somewhat brief history about myself, and I hope it helps in any kind of way - by increasing insight, learning something, etc. |
| not_optimism | olanzapine | HOW DARE YOU? This is a voice from within, spoken so loud it shakes me, and I projectile vomit the contents of my stomach, watery acidic technocolor psychedelic yawn, I am crying. HOW THE FUCK DARE YOU VIOLATE THE HOLY? |
| pride | amphetamine | I got her some water and told her I know how she feels. I felt so proud of myself for turning it down, and without seeing the consequences of messing up my life I would have never been that strong. |
| not_pride | haloperidol | A sizeable number of audience members had apparently asked the network to conceal their identities (lol) so their faces were blurred out; the collective effect of these things was, at the time, incomprehensibly disturbing. |
| realization | lsd | All of the repeating patterns I had observed in nature made me realize that things repeat over and over again. I thought about how much Fleur looked like my first girlfriend, over 20 years ago, and realized that there were only so many ways to make hands, arms, a face. |
| not_realization | mirtazapine | Good luck and happy tripping everyone. |
| relief | mdma | N held all the pills, so I wasn't worried about security. After a quick pat down, we were free of the crowd and the temperature dropped by 15 degrees. It was a relief to be able to move freely again and my mood slowly improved. |
| not_relief | mirtazapine | Why is it a chair? What makes a chair a chair? Where did the idea of a chair come from? Chair, that’s a funny word.” The processes of my mind could not keep up with all the questions I had and everything I wanted to know. |
| remorse | hydrocodone | I essentially felt nothing other than the DRUG high, although I did feel very guilty for my outburst previously in the evening. I apologized profusely in the car and blamed it on the Vicodin. |
| not_remorse | risperidone | My friend and I made our way to a secluded spot in a this wildreness park. The spot was cool, totally covered by trees, there were tons of branches intertwined in a dome shape, and a log for sitting on. |
| sadness | lsd | I pick up my glasses to put them on and the asymmetry of my head is enormous. I feel my teeth grind against each other, feel how much my body spirals, and I'm sad. |
| not_sadness | haloperidol | She listened to every word that I had to say, and nodded with understanding. I marveled at how well we were communicating. Whenever I spoke I knew that I had her full attention, and in turn, when she spoke I listened to everything that she had to say. |
| surprise | lsd | I was shocked. I asked her, “So this was all a trip?” She told me that it was. I was shocked and almost disbelieving. I couldn’t believe that a simple drug had been the culprit of this madness. |
| not_surprise | 2c-d | We had the following selection: Tool, A Perfect Circle, Smashing Pumpkins, Appleseed Cast, and some other prog rock. For anyone unfamilar with the musical experience on MDA,I recommend choosing anything that you like, don't get stuck with anything too abrasive or annoying. |
| neutral | morphine | -high timer |
| not_neutral | haloperidol | Thank you. Thank you. I love you. Love and Blessings Always and Forever. |

#### Supplementary Table 2: Extreme BERTowid

Below are the highest and lowest ranking ~32 word testimonial-windows for each of the 35 tags learned by BERTowid. The precision with which each tag matches the meaning for many of these passages flagged by BERTowid–and also their opposites–suggest that these descriptions are meaningful in both the positive and negative directions for several dimensions of analysis.

| tag | drug | text |
| --- | --- | --- |
| Small_Group | psilocin | I think it will be helpful if you are contemplating trying DRUG or are just reading about other peoples' experiences. Here we go... P and I were both 16 years old at the time and had never done shrooms. |
| Not_Small_Group | 2c-i | I woke up on the third day of a four-day music festival, slightly hungover and groggy from a batch of DRUG I had taken the previous morning. After a brief wakeup period going through the motions of dressing, tooth brushing, etc, I decided to finish off a batch of DRUG I had been sold the previous day. |
| General | ketamine | Good thing I didn’t DRUG it with that dirt. I located the muscle (vastus lateralis) and relaxed it (DON’T flex it while injecting). The site was cleaned with DRUG and I found my self sitting with a DRUG ready to inject. |
| Not_General | cocaine | It was a temporary fix. It changed nothing about me. It did not improve the quality of my life. To put it simply, it was a fake, empty experience. It was a temporary fix. |
| First_Times | cocaine | So before long a guy friend of mine asked if I was interested in trying some DRUG I said sure. Let me give y'all a little more background on myself. I'm a staight A student, I'm on the honor roll, I'm on the debate team, and in many clubs at school. |
| Not_First_Times | lsd | Indulging in one of these sins often leads to you indulging in another, then another, then another, and eventually all of them. All these things have a web effect. Problems in one area resulted in problems in others. |
| Alone | psilocin | I stood up and opened my closet, withdrew my DRUG pipe and consumed deeply. I sat down in the dark again, I felt better, though now I decided I better take this lying down. |
| Not_Alone | 2c-e | Me and my boyfriend both were shaking and we were bouncing our legs and our legs started shaking so fast it was like they were machines we were just witnessing shake like mad. The shaking almost felt involuntary because if we didn't shake everything was just painful. |
| Difficult_Experiences | dxm | He asked me 'Why did we do this?', and I could not answer. He then said 'This is the worst experience of my life'. I was thinking something along the lines of 'Oh it's not THAT bad. |
| Not_Difficult_Experiences | mdma | and doing as much research on E and all its permutations, effects... I started a daily program of DRUG in the morning, with Gingko Biloba or St. John's Wort, followed by DRUG and Valerian at night before bed. |
| Glowing_Experiences | ibogaine | DRUG DRUG Married, female, age 42. 115 pounds 19 months detoxed from DRUG muscle relaxants & DRUG anti-depressants DRUG Fanatic since August of 2009. Longest trip of my life to date - 14-16 hours Ever since I “broke my head” in December, 2009 with a series of extremely rough DRUG breakthroughs, I have had this nagging and oppressive thought that my death is near, sometimes imminent. |
| Not_Glowing_Experiences | dxm | I don’t remember all that well, but I got myself into the same type of situation. I figured, hey, I’ve dropped DRUG a few times, I can handle a DRUG binge. Well, all that I really remember was being attacked and defeated by a demon by the name of Rez. |
| Retrospective_Summary | mdma | To put recent events into perspective, my DRUG history started off with a bang eight years ago at the age of 21. I continued experimenting with it once or twice a month for the next six months culminating in the realization that the destructive effects were increasing and overtaking the beneficial effects with each new visit. |
| Not_Retrospective_Summary | diphenhydramine | I had parked my car half-way on the sidewalk with the windows down at 3am. It was only 3 blocks from where I had ended up. My money was miraculously still there, sitting all over the seat. |
| Various | 2c-i | At around 5 am, I was walking around my house, and I started noticing that the place was an absolute mess, which never seems to really touch me when I have people over, but that day it did. |
| Not_Various | methamphetamine | There is no accompanying orchestral orgasm of consciousness, you just suddenly realize that you've accidentally crept into this new dimensional level of consciousness. And it is here that you will experience The Others, though we will refrain from commenting on this aspect until the need for subsequent missives on the subject arises. |
| Unknown_Context | 5-meo-tmt | Mild nausea about 1 and a half hours after dosing. Nausea lasts for about 2 hours and is temporarily relieved when gases leave the body. After the nausea passes, the familiar stomach feeling of DRUG sets in - not quite nausea, just an indescribable odd feeling that is easily ignored. |
| Not_Unknown_Context | ketamine | DRUG DRUG DRUG DRUG Background: Healthy, casual psychonaut. Average diet and exercise. Prior familiarity with a wide range of psychoactives spanning ten years of use. Set/Setting: I'm in my apartment with one of my roommates. |
| Mystical_Experiences | psilocin | I lost myself in the ebb and flow of the universe and I was home. I could go on for hours and hours, and have before, about the wonders of my awakening, but truly, words don't exist for what I was experiencing, so hopefully |

| | | |
|---|---|---|
| | | my meager attempt at explanation will do. |
| Not_Mystical_Experiences | methamphetamine | I started attendin NA, Narcotics Anomynous, meetings 4 or 5 times a week. My life today is DRUG free. I havent touched it in a little over 2 months. I'm alot happier off of it. |
| Health_Problems | thc | I then consumed 3 more the next morning, and that was it. This was about 6 weeks before the experience I am writing about here. other then this I have had local anaesthetic twice, once for repairing a broken tooth, and once for stitches to a cut. |
| Not_Health_Problems | 2c-e | I was unconcerned about any other humans, I felt that I was alone in this new dimension and yet I was in awe and mostly unafraid. My destination was the park, a rather large one comprising 3 open fields, trees on the border and some children's play equipment. |
| Combinations | dmt | No problems. |
| Not_Combinations | psilocin | The walls and ceiling were breathing in and out together to a great degree, and everything on my wall was 3-D. People came out of the wall, and a space shuttle was in constant flight, flying past stars and galaxies, in a poster where it is just taking off. |
| Not_Applicable | cocaine | The first binge lasted 2 or 3 months within the first week of using I lost 15 pounds, I thought I found my answer to everything in life just a simple line, a quick fix and everything was all right. |
| Not_Not_Applicable | mdma | He was completely sober at the time, but had an 18 of beer in his trunk and we had 5 e pills along with some DRUG a few bowls. We just kept calm and explained it off, and the cop just told us to get where we were going and stay there. |
| Bad_Trips | lsd | All my lives were the same from beginning to end. I did the same things and got the same ending. Looking back at it, I realize that it was true insanity that I experienced in my bad trip. |
| Not_Bad_Trips | 5-meo-dmt | At the beginning of the ceremony we were given the space to request what it is we wanted from the medicine. I didn't take this lightly, most of my group and many others got exactly what we asked for and to a major degree. |
| Hangover_Days_After | 2c-b | returning home. |
| Not_Hangover_Days_After | thc | It is quite useful actually, that I am able to see all the keys and the screen at the same time. God, those fingers are moving fast and make a fun little dance on the keyboard! |
| Entities_Beings | mescaline | Closed eyed visuals are beyond amazing. I seem to have met a Peruvian shaman and my body/soul is filled with brilliant purified white light. I am seeing closed eye visuals of the shaman and then the vision switches to someone who seems to be in Tibet. |
| Not_Entities_Beings | alprazolam | At about T+45 began to feel the effects of the drug. Mostly a mild stoning effect, but motion seemed incredibly interesting. I left the group I was with (always a bad idea when on a new substance) and went to hang out with some of my other friends who were drinking. |
| Music_Discussion | 5-meo-dmt | Life is good. |
| Not_Music_Discussion | oxycodone | (meaningless term unless youre a shooter I know) It resembled mud, but I managed to get the shit in the DRUG after some work. POKE, ASPIRATE, BURNING VEINS, ENDORPHIN EXPLOSION, BLISS, COLLAPSE. Thats the jist of it, but I lived. |
| Addiction_Habituation | methamphetamine | I will always have a problem coping with my problem. I went through so much pain and suffering (not to mention the pain I put my family through) due to my addiction. |
| Not_Addiction_Habituation | cocaine | I was ready to take a little walk alone and after getting dressed followed the advice and took a couple of big bumps off a key. I chilled out for a few minutes expecting something insanely euphoric and almost spiritual. |
| Post_Trip_Problems | Salvinorin A | I took the third hit, holding all of it deep in my lungs. Then NOTHING. I lost consciousness. What I gathered from the first hand account of friends was that I stumbled around very clumsily. |
| Not_Post_Trip_Problems | tma-2 | Back home (0:25 pm ie I checked my pupil and it was still dilated although only moderately so. I had expected to be baseline after 8h, so occupied myself until 1:40 am expecting to be out of the experience at that point. |
| Nature_Outdoors | lsd | Straight ahead was the DRUG clear ocean, sandy beaches and creamy white clouds of the 'east-side,' while directly behind us was the immense green landscapes of the thick Hawaiian jungles and it's dark, looming rain clouds. |
| Not_Nature_Outdoors | amphetamine | The sad thing is though, I like DRUG ALOT. And even though this happened to me today, I don't know if I'm going to have enough will power to stop taking it. It's not as if the drug is addictive in such a way that your body feels it needs it to function, it's more like a mental drug. |

| | | |
|---|---|---|
| Relationships | ibogaine | My family have lived for generations in the Caribbean, unwittingly participating in - yet outwardly shunning - the subversive caste system. I accepted it unquestioningly, at home being an unspecified outcast in my society, marked by my skin and my eyes, my base state subtly asserted as that of an ‘oppressor, an outsider’ while in truth my childhood identified me as the oppressed. |
| Not_Relationships | thc | - But then-- ALL thoughts are hallucinations! At that point I would convulse involuntarily. From somewhere deeper in my brain than logic, I saw an infinite loop of thoughts stretching into darkness ('Then that statement, too, must be a hallucination! |
| Depression | mdma | DRUG DRUG DRUG For years I've experienced various levels of depression (3 years). For the first 2 years I was diagnosed with major depression after two attempts at my own life. I sought treatment and was put on zoloft. |
| Not_Depression | dmt | They arent making it up, it is really there. It just doesnt have the same dimensions as this 'real' world. The efficiency of there seemed immaculate. Everything was in perfect time/tune and there was, dare I say it? |
| Therapeutic_Intent_or_Outcome | lsd | DRUG catalyzed the first major shifts in my spinal structure and gave me the experience of a straight spine, providing me with hope, encouragement, and wisdom that strongly impacts my mental experience of my body in everyday life. |
| Not_Therapeutic_Intent_or_Outcome | 5-meo-mipt | Very nice... I can see how some might compare this to the DRUG DRUG I feel that listening to some music, laid out, with my eyes closed would be pristine. I doubt it'll happen though. |
| Overdose | 25i-nbome | I'm going to die anyways, so why let myself suffer?' I knew how I would have killed myself, too- I was going to get out of Chayla's house and lay in the street and wait for a car to hit me. |
| Not_Overdose | 2c-i | This instantly brings some peace of mind back, I can feel the soft cradle of DRUG rock my hopped up mind softly back and forth. I feel secure in my abilty to discern between what is real and what is not. |
| Medical_Use | methadone | I had no time for a girlfriend, no time for my family, and definately no time to goto school even though it was my senior year. I tried an inpatient 30 day shit but that just put my problem on hold until I got out and then went into an IOP (Intensive Outpatient Program) that lasted a minimum of six months. |
| Not_Medical_Use | methamphetamine | So I tried it. I finally smoked some DRUG I took one hit of some good quality glass. It wasn't a large hit, I didn't feel much at all. I took another and began to feel alert and awake, as I expected. |
| Sex_Discussion | amphetamine | But be weary, shes a tempting mistress, who could possess you in a matter of weeks.* |
| Not_Sex_Discussion | methamphetamine | I was not overly keen to try it, as I tend to mainly enjoy a monthly or two-monthly dose of DRUG and don't feel I need any other drugs, having been down the psychodelics path over many years and becoming well and truly over them. |
| Train_Wrecks_Trip_Disasters | ketamine | I fell asleep soon afterwards. |
| Not_Train_Wrecks_Trip_Disasters | mescaline | The feeling of the DRUG is swimming happily through my body, a school of fish frolicking about, miles beneath the surface. Writing is flowing effortlessly and readily from mind to page. Very little editing is being done at this point. |
| Guides_Sitters | lsd | From the MAPS Bulletin - Volume 9 Number 2 Summer 1999 - pp. 11-14 ------------------------------------------------------------------------ Successful Outcome of a Single DRUG Treatment in a Chronically Dysfunctional Man Gary Fisher, Ph.D. ------------------------------------------------------------------------ Editor's Note: MAPS publishes accounts like the one below because they are reminders of the sometimes surprising powerful effects that psychedelics can have. |
| Not_Guides_Sitters | methamphetamine | Don’t expect your will to be greater than the will of DRUG It has pulled me in countless times despite negative consequences. I plan to quit using altogether soon however I am indulging tonight. |
| Rave_Dance_Event | mdma | The idea of such a drug disgusted me. However we went to a rave the following night and I dropped a mitsubishi. That was my first rave and probably my best. We were dancing in some guy's apartment until the early hours of the morning. |
| Not_Rave_Dance_Event | lsd | I love lysergic DRUG diethylamide, it is an unbelievably wonderful sacrament that has been bestowed upon the human race, and I truly believe that a direct experience of reality, beyond labels or concepts, could push the human race off the destructive path it is currently on. |
| Preparation_Recipes | morphine | *If you are not using plant material, skip this step* Both stems and pods can be used for their alkaloids, but being plants the quality is variable. It is best to harvest shortly after the petals fall from the plant and before the plant is flushed of its goodies become lost from the rain. |
| Not_Preparation_Recipes | mescaline | I speak to someone they tell me I'm five mile from my home. I believe I can make. I get back on the road. I feel like I am losing it. |

| | | |
|---|---|---|
| Festival_Lg_Crowd | doi | Walking into the sea of people was an intense and disorienting experience, ACL is a big festival, there were thousands and thousands of people there all at the main stage, because almost everyone wanted to see Dylan. |
| Not_Festival_Lg_Crowd | methadone | After struggling slowly facing one thing at a time I managed to get my license back and got a car (nothing fancy I am saving for a corvette or the new camaro). I have a job as the senior chemist making serious money which I have managed to save a little of for emergencies. |
| Health_Benefits | methadone | First off, a little bit about myself. I have been using DRUG and opioids for 2 years non-stop to deal with pain resulting from a kidney condition. I've taken DRUG DRUG meperidine, DRUG propoxyphene, DRUG and now finally methadone. |
| Not_Health_Benefits | ketamine | So, no real worries going in. I was expecting something fun like mxe, especially since I had always heard of it in the dance and rave contexts. I was at my friend E's grandparent's house (they were gone and had left him the keys) and we were gonna try the DRUG recently procured. |
| Large_Group | dxm | and good night. |
| Not_Large_Group | venlafaxine | Having experienced both, I asked my new, smart, shrink. He said that the effect is a severe defficiency of serotonin as well as neurepinephrine, because the stimulus to produce more is gone and the brain hasn't gone back to producing enough of its own yet. |
| Multi-Day_Experience | dob | I pay him ten for two hits- half what he was asking- and promise that if and when I start to feel it I'll pay him the other ten. I stick em under my tounge and proceed to the show to beat people up for the next two hours. |
| Not_Multi-Day_Experience | lsd | Here comes five pairs of headlights! Yeah, the cab is here. We walked towards the van, waving it down. When it stops, we realize it wasn't a cab, it was a mom with a carload of kids inside. |
| Club_Bar | 5-meo-dipt | The buzz increased a fair bit, and probably would have probably resulted in a pleasant experience had we not been in the very hot, crowded, noisy nightclub environment (it was Easter weekend so the club was unusually packed). |
| Not_Club_Bar | 5-meo-dmt | (days later->) Now, This is where my tryps get interesting, Things that shouldn't happen, do. I have 2 digital clocks in my room, One with the correct time, and the other I plugged in 10 minutes after midnight, so that it would always flash the near correct time. |

### Supplementary Table 3: Drug Information

| Pharmacological Class | Drug Name | Common and Brand Names | Chemical Class | Non-Medical Dosage Range (in milligrams) | Therapeutic Dosage Range (in milligrams) | Average Duration of Action (in hours) | Route(s) of Administration |
|---|---|---|---|---|---|---|---|
| Stimulant | **Amphetamine** | *Speed, Adderall, Pep* | phenethylamine | 2.5-50 | 5-40 | 5 | Oral, insufflation |
| | **Cocaine** | *Coke, Coca, Crack, Blow, Girl, White, Snow, Nose Candy* | tropane alkaloid | 5-90 | - | 0.82 | Insufflation, IV, oral |
| | **Methamphetamine** | *Meth, Speed, Ice, Glass, Shard, Tina, Crank, Desoxyn* | amphetamine | 5-60 | 20-60 | 9.5 | Oral, insufflation, IV |
| Psychedelic | **25i-NBOME** | - | phenethylamine | 1-16 | - | 15 | Oral |
| | **2C-B** | *Nexus, Bees* | phenethylamine | 5-45 | - | 6.5 | Oral |
| | **2C-C** | - | phenethylamine | 5-90 | - | 6 | Oral |
| | **2C-D** | *2C-M, LE-25* | phenethylamine | 3-00 | - | 4 | Oral |
| | **2C-E** | *Eternity, Aquarust* | phenethylamine | 2-30 | - | 8 | Oral |
| | **2C-I** | - | phenethylamine | 2-30 | - | 8 | Oral |
| | **2C-P** | - | phenethylamine | 3-30 | - | 8 | Oral |
| | **2C-T-2** | *Rosy* | phenethylamine | 3-30 | - | 8 | Oral |
| | **2C-T-4** | - | phenethylamine | 8-20 | - | 15 | Oral |
| | **2C-T-7** | *Beautiful, Blue Mystic, 7th Heaven* | phenethylamine | 3-40 | - | 8 | Oral |
| | **5-MeO-DiPT** | *Foxy Methoxy, Foxy* | tryptamine | 3-20 | - | 6 | Oral |
| | **5-MeO-DMT** | *The God Molecule, Toad, Jaguar, Soma* | tryptamine | 1-20 | - | 0.5 | Oral |
| | **5-MeO-MiPT** | *Moxy* | tryptamine | 3-20 | - | 6.5 | Oral |
| | **5-MeO-TMT** | - | tryptamine | 75-150 | - | 7.5 | Oral |
| | **DiPT** | - | tryptamine | 15-150 | - | 4.5 | Oral |
| | **DMT** | *N,N-DMT, Dmitry, The Glory, The Spirit Molecule* | tryptamine | 2-60 | - | 0.2 | Inhalation |
| | **DOB** | *Bromamfetamine, Bromo-DMA* | amphetamine | 0.2-3 | - | 19 | Oral |
| | **DOI** | - | amphetamine | 0.5-3 | - | 20 | Oral |
| | **DOM** | *STP (Serenity, Tranquility, and Peace)* | amphetamine | 0.5-10 | - | 14 | Oral |
| | **DPT** | *The Light* | tryptamine | 50-350 | - | 3 | Oral |
| | **Ibogaine** | *Endabuse, Iboga* | tryptamine | 15 | - | 45 | Oral |
| | **LSD** | *LSD-25, Lucy, L, Acid, Cid, Tabs, Blotter* | lysergamide | 0.015-0.3 | - | 10 | Oral |
| | **Mescaline** | *Mescaline, Peyote, San Pedro, Cactus, Buttons, Devil'* | phenethylamine | 50-800 | - | 11 | Oral |
| | **Psilocin** | *Psilocin, Psilocine, Psilocyn, Psilotsin* | tryptamine | 5-40 | 20-30 | 5 | Oral |
| | **TMA-2** | - | amphetamine | 5-60 | - | 10 | Oral |
| Opioid | **Hydrocodone** | *Vicodin, Zohyrdo ER* | morphinan | 3-40 | 10-80 | 6 | Oral, insufflation, IV |
| | **Hydromorphone** | *Dilaudid* | morphinan | 0.5-8 | 1-12 | 3.5 | Oral, insufflation, IV |
| | **Methadone** | *Dolophine* | diphenylpropylamine | 1-30 | 10.0-120 | 6 | Oral |
| | **Morphine** | *MSContin, Oramorph, Zomorph, Sevredol* | morphinan | 10-30 | 1-10 | 5 | Oral, insufflation, IV |
| | **Oxycodone** | *OxyContin, Oxy, Roxicodone, Oxecta, OxyIR, Endone, Codilek, Redocam* | morphinan | 1-40 | 2.5-40 | 5 | Oral, insufflation, IV |
| Hallucinogen | **Salvinorin A** | *Salvia, Salvia Divinorum, Diviner's Sage, Ska Maria Pastora, Seer's Sage, Sally* | salvinorin | 0.2-1.0 | - | 0.9 | Smoked |
| | **THC** | *Cannabis, Marijuana, Weed, Pot, Mary Jane* | cannabinoid | 0.4-10 | 2-20 | 3.65 | Smoked, oral |
| Entactogen | **MDA** | *Sass, Sassafras, Tenamfetamine* | amphetamine | 20-145 | - | 6.5 | Oral |
| | **MDMA** | *Molly, Mandy, Emma, MD, Ecstasy, E, X, XTC, Rolls, Beans* | amphetamine | 10.0-180 | 80-120 | 4.5 | Oral |
| Dissociative | **DXM** | *DXM, DMO, DM, Dex, Robitussin, Delsym, DexAlone, Duract* | morphinan | 10.0-700 | 10-60 | 10 | Oral |
| | **Ketamine** | *K, Ket, Kitty, Special K, Cat Tranquilizer, Ketaset, Ketalar, Vitamin K, Purple, Jet* | arylcyclohexylamine | 5.0-1000 | 25-100 | 1.25 | IV, IM, buccal |
| | **PCP** | *PCP, Angel Dust, Sherman, Sernyl, Wet, Dust, Supergrass, Boat, Tic Tac, Zoom* | arylcyclohexylamine | 1.0-15 | - | 6 | Smoked, oral |
| Deliriant | **Diphenhydramine** | *Benadryl* | ethanolamine | 25-700 | 25-50 | 6.5 | Oral, IM, IV |
| | **Mirtazapine** | *Avanza, Axit, Mirtaz, Mirtazon, Remeron, Zispin* | piperazinoazepine | 3.5-250 | 15-60 | 18 | Oral |
| Depressant | **Alprazolam** | *Xanax* | benzodiazepine | 0.1-3 | 0.25-2 | 6.5 | Oral, IV |
| Antipsychotic | **Aripiprazole** | *Abilify* | piperazine | - | 2-30 | - | Oral, IM, LAI |
| | **Haloperidol** | *Haldol* | butyrophenone | 0.25-10 | 2-30 | 24 | Oral, IM, LAI |
| | **Olanzapine** | *Zyprexa* | thienobenzodiazepine | - | 2.5-40 | - | Oral, IM, LAI |
| | **Quetiapine** | *Seroquel, Xeroquel, Ketipinor* | dibenzothiazepine | 10.0-200 | 25-800 | 6 | Oral |
| | **Risperidone** | *Risperdal* | benzisoxazole | 0.25-6 | 0.25-6 | 16 | Oral, IM, LAI |
| Antidepressant | **Bupropion** | *Wellbutrin, Zyban, Aplenzin* | aminoketone | 75-325 | 100-450 | 10 | Oral |
| | **Paroxetine** | *Paxil, Seroxat, Loxamine* | piperidine | - | 12.5-75 | - | Oral |
| | **Sertraline** | *Zoloft* | SSRI | - | 25-200 | - | Oral |
| | **Venlafaxine** | *Effexor* | SNRI | - | 75-375 | - | Oral |
